# Neprilysin mediated cleavage of phospholamban dysregulates SERCA in heart failure

**DOI:** 10.64898/2026.06.23.732949

**Authors:** Jacob D. Cunningham, Taylor A. Phillips, Alexis R. Mazzenga, Kush N. Nagrani, Tuan H. Bui, Seby Edassery, David Y. Barefield, Seth L. Robia

## Abstract

**Background:** Neprilysin (NEP) is a zinc-dependent metalloprotease targeted in heart failure therapy to prevent it degrading circulating cardioprotective vasoactive peptides. NEP can also cleave sarcolipin (SLN), the skeletal- and atrial muscle-specific micropeptide regulator of the sarcoplasmic reticulum Ca^2+^-ATPase (SERCA). A direct pathophysiological role of NEP in ventricular muscle has not been established.

**Methods:** Proteomics and immunoblot analysis of human myocardial specimens were used to quantify NEP abundance in failing and non-failing hearts. Heterologous protein expression and biochemical binding assays assessed NEP-mediated cleavage of phospholamban (PLB) and its impact on PLB–SERCA interactions. Functional consequences of NEP expression or inhibition were evaluated in neonatal rat ventricular myocytes and in a human induced pluripotent stem cell–derived cardiomyocyte (hiPSC-CM) model of heart failure.

**Results:** We observed increased NEP abundance in failing human myocardium relative to non-failing controls. We demonstrated that NEP cleaves phospholamban (PLB), disrupting PLB–SERCA interactions. Mutation of PLB (V49A), prevented NEP cleavage and preserved PLB-SERCA binding, indicating V49 is critical for NEP substrate recognition. In neonatal rat ventricular myocytes, NEP expression was associated with faster Ca^2+^ transient decay kinetics and increased SR Ca^2+^ load, consistent with reduced SERCA inhibition. Inhibition of NEP in a hiPSC-CM heart failure model attenuated the hypertrophic transcriptional responses and reversed Ca^2+^-transport dysregulation.

**Conclusions:** These findings implicate increased NEP expression in the sarcoplasmic reticulum of cardiomyocytes as previously unrecognized maladaptive consequence of heart failure contributing to cardiac dysfunction. In this novel pathophysiological mechanism, increased NEP results in PLB cleavage and loss of regulation of SERCA. While this may relieve SERCA inhibition and augment cellular Ca^2+^ handling, loss of PLB chronically disrupts heart’s dynamic response to adrenergic stress, changing heart rate, or other physiological challenges. The data provide new insight into the cardioprotective effects of pharmacological NEP inhibition in clinical practice, reveal a novel mechanism of action of neprilysin inhibition in cardiomyocytes and may help inform future therapeutic strategies for patients with heart failure.

**Graphical Abstract:** 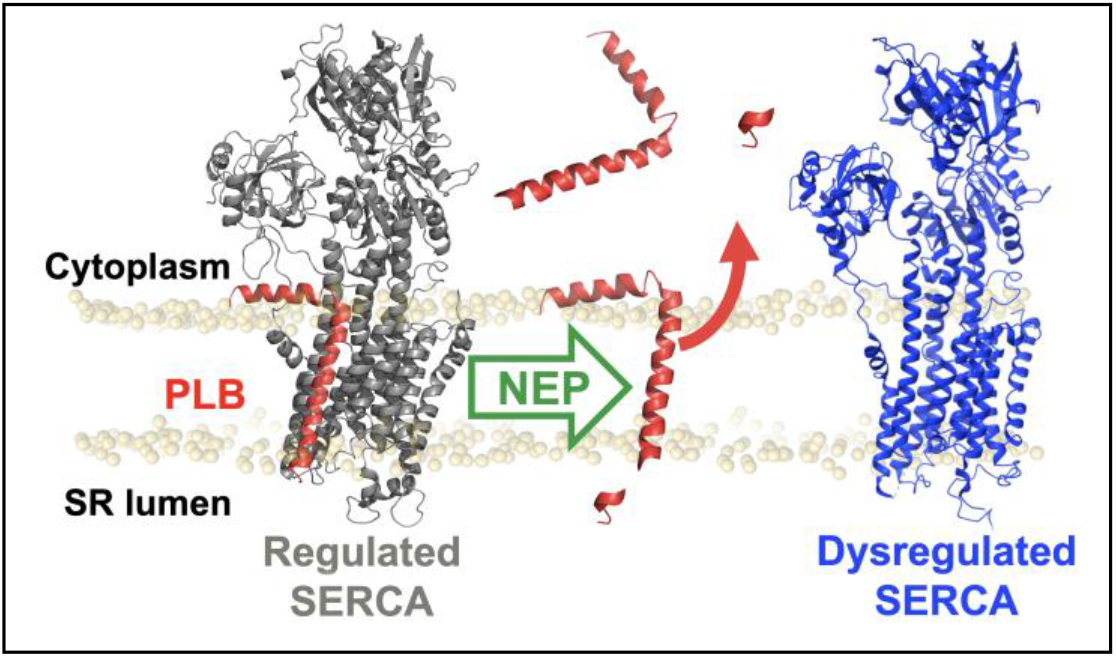

## Introduction

Neprilysin (NEP) is a broadly-specific, zinc-dependent metallopeptidase with more than 50 reported substrates^1-4^. It is expressed in the heart, kidneys, brain and vasculature where it is typically anchored to the extracellular side of cell plasma membranes, but it can also circulate throughout the body after being released from the cell by cleavage of its extracellular domain^5^. Neprilysin inhibition using angiotensin receptor–neprilysin inhibitors (ARNIs) has emerged as one of the four cornerstone therapies for patients with heart failure (HF) with reduced ejection fraction (HFrEF), reducing overall mortality by ∼30%^6^. Furthermore, recent trials have suggested that ARNIs may be beneficial for certain patients with HF with preserved ejection fraction (HFpEF)^6,7^. These clinical benefits are generally attributed to preservation of circulating vasoactive peptides like ANP and BNP that would otherwise be degraded by NEP^2^. In addition to this canonical role as an extracellular peptidase, recent evidence revealed a catalytically active pool of membrane-bound NEP localized to the sarcoplasmic reticulum (SR)^8^, suggesting NEP may have cardiomyocyte specific substrates^1,7,9-12^. Notably, NEP is significantly increased in the hearts of patients with dilated and hypertrophic cardiomyopathy^11^. Together, these findings raise the possibility that NEP may cleave proteins involved in cardiomyocyte function and contribute to HF pathogenesis.

Recent work by Schiemann et al. demonstrated that NEP cleaves sarcolipin (SLN), a micropeptide that regulates the sarco(endo)plasmic reticulum Ca^2^+-ATPase (SERCA) in skeletal and atrial muscle^13,14^. NEP cleavage removes five C-terminal residues of SLN that are functionally important for inhibition of SERCA. In ventricular cardiomyocytes, the predominant SERCA regulatory micropeptide is phospholamban (PLB). Decades of structural and functional studies have shown SLN and PLB regulate SERCA through similar mechanisms^15-17^. Given these similarities, we hypothesized that PLB could also be a substrate of NEP in the heart. Like SLN, PLB binds to SERCA and slows SR calcium reuptake by reducing the pump’s apparent affinity for Ca^2+^ under basal conditions. During β-adrenergic stimulation, phosphorylation of PLB relieves this inhibition, increasing SERCA activity and enhancing myocardial relaxation^18^, allowing the heart to dynamically respond to exercise and adrenergic stress^19,20^. Disruption of this regulatory mechanism has been implicated in dilated cardiomyopathy, arrhythmogenesis, and impaired calcium handling^21-23^. Our prior work demonstrated even small deletions within the C-terminal transmembrane helix of PLB can abolish its inhibitory potency^24^. Thus, NEP removal of C-terminal residues from PLB, as it does from SLN, could significantly disrupt ventricular Ca^2+^ handling. Therefore, pharmacological inhibition of NEP by ARNIs could alter cardiac function directly by sparing PLB and preserving normal SERCA regulation. Studies of PLB expression changes in HF have been inconsistent^3,4,25,26^, but many suggest decreased PLB protein levels in HF^27-29^. Consistent with a possible role for NEP in proteolysis of PLB, our recent quantitative mass spectrometry study showed a reduction in membrane-associated PLB^30^.

Here we investigated whether aberrant NEP protease activity causes maladaptive changes in the failing heart. We tested whether NEP expression is altered in failing human myocardium and whether it can cleave PLB to modify SERCA function. Our results suggest that NEP contributes to calcium mishandling in HF, which may account for some of the cardioprotective effect of ARNI therapy.

## Results

### Neprilysin is increased in human dilated cardiomyopathy

Previous studies have shown that the gene transcripts of neprilysin (NEP, *MME*) are increased in dilated cardiomyopathy and hypertrophic cardiomyopathy^11^. Here, we performed mass spectrometry to evaluate whether the protein levels of NEP are increased in human heart failure (HF). Membrane-associated proteins like NEP are often underrepresented in standard LC-MS/MS workflows^31,32^ because membrane-embedded sequences are poorly soluble in aqueous buffers^33,34^, require detergents for extraction, and yield large hydrophobic tryptic peptides due to the relative paucity of Lys/Arg cleavage sites within transmembrane domains. Thus, membrane peptides often require detergent solubilization, which can further suppress ionization^34^. To improve detection of membrane-associated species for the present study, tissue from the left ventricle of human hearts was subjected to membrane enrichment via differential ultracentrifugation as described previously^35^, followed by delipidification^33^, digestion with trypsin/LysC, and mass spectrometry (MS) (**Figure 1A**). Using human myocardial samples from failing hearts and non-failing rejected donor controls, we performed differential proteomic analysis to define protein-level remodeling in HF. Donor and patient characteristics, including age, sex, and available clinical metrics are summarized in **Supplementary Table 1**. Applying thresholds of adjusted *p* < 0.05 and |log_2_ fold change| ≥ 1 identified 346 significantly altered proteins in HF compared with NF controls, including 186 increased and 160 decreased proteins (**Figure 1B**). Among the proteins found to be elevated in HF samples, NEP exhibited a 4.1-fold increase in HF samples compared with NF controls (**Figure 1C**). Western blot analysis showed a 3.4-fold increase of NEP in HF hearts relative to NF hearts (**Figure 1D, E**), consistent with proteomic analysis. To determine the cellular and subcellular distribution of NEP in cardiomyocytes, we performed immunofluorescence microscopy of NF and HF human cardiomyocytes, comparing the localization of SERCA (blue) and NEP (green) with phalloidin-labeled F-actin (red) (**Figure 1F**). NEP was markedly elevated in HF myocytes compared with NF, consistent with proteomics and Western blot data. NEP exhibited a punctate distribution, with enrichment at cell-cell junctional regions (intercalated discs) (**Figure 1F, Region 1, arrow**). The relative enrichment of NEP at the intercalated disc is presumably due to canonical plasma membrane localization of extracellularly-facing NEP and the high membrane content of this subcellular region. Adjacent cell membranes interdigitate at the intercalated disc, providing a large contact area for cell-cell communication. Thus, any plasma-membrane-localized protein may be expected to be relatively concentrated there. We also observed a fraction of intracellularly-localized NEP, which was arranged in longitudinal streaks that ran parallel to the sarcomere axis (**Figure 1F, Region 2**). Notably, this NEP signal coincided with SERCA in voids between myofibrils (**Figure 1F, Region 2, arrows**), consistent with localization in the SR membrane. The significance of the punctate pattern for NEP localized in the plasma membrane and SR membranes is unknown. We considered that this pattern could be an artifact of IF localization, but we did not observe significant puncta in control experiments in which we applied secondary antibody alone (**Figure S1**). Moreover, the number of puncta was markedly increased in HF vs. NF, so we conclude that this signal represents a bona fide NEP localization pattern. We also observed strong colocalization of NEP, PLB, and SERCA tagged with fluorescent proteins and expressed in HEK293T cells (**Figure S2**).

**Figure 1.**
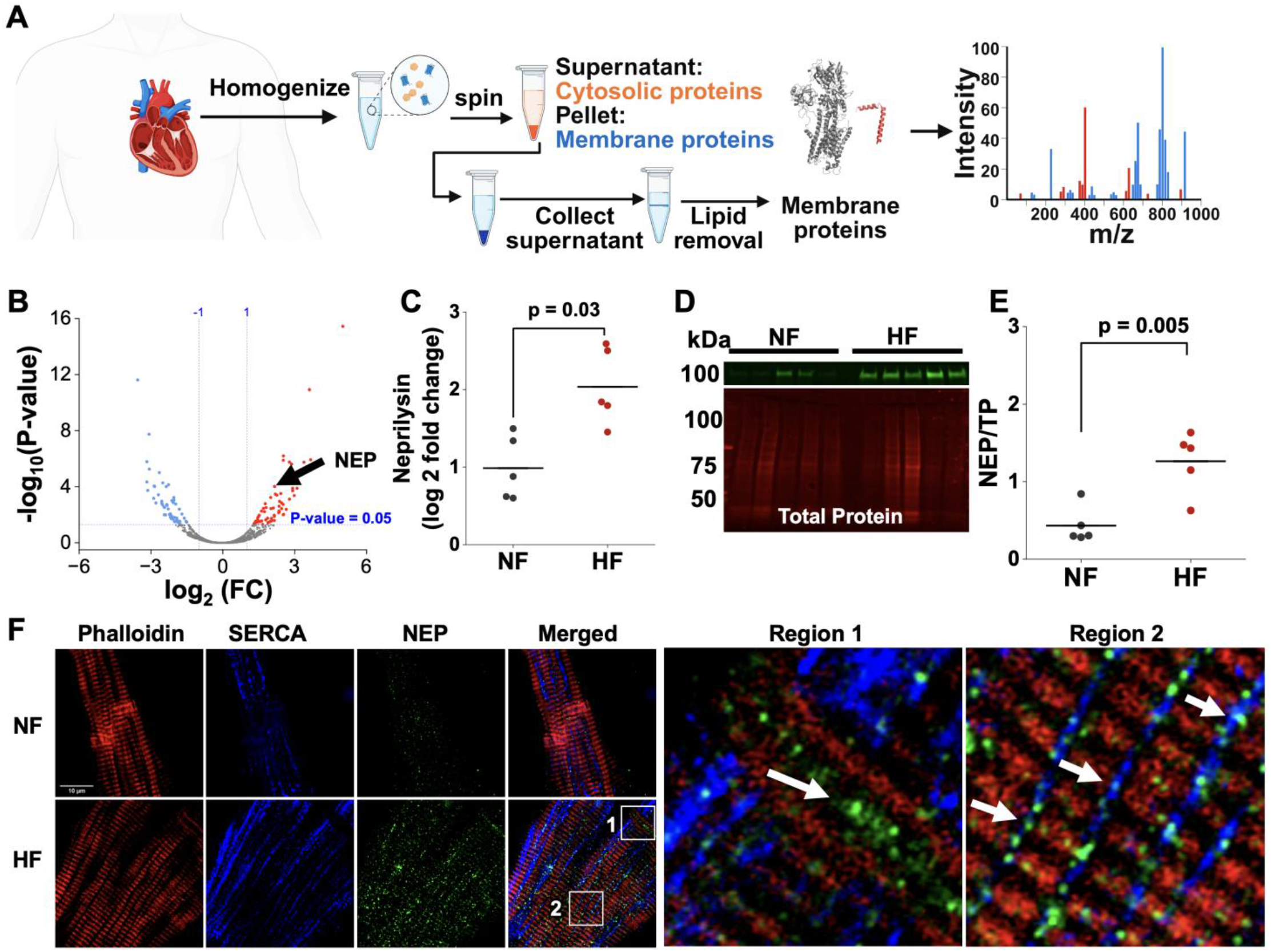
**A)** Schematic of human heart tissue membrane protein enrichment and delipidification workflow prior to mass spectrometry. **B)** MS results of differentially expressed proteins in HF versus NF comparison. **C)** Quantification from the mass spectrometry data specifically for NEP (gene name: MME). **D)** Western blot for NEP for non-failing (NF) and heart failure (HF) left ventricle samples. **E)** Summary data from data in **C** (N=5). p values were calculated via t-test or adjusted t-test. **F)** Immunofluorescence microscopy of NF and HF human ventricular myocytes revealed neprilysin (green), SERCA (blue), and actin (red). Boxes 1, 2 in the HF, Merged image are enlarged at right. Arrows indicate NEP localized at the intercalated disc (**Region 1**) or overlapping with SR-localized SERCA in intermyofibrillar regions (**Region 2**) of failing human cardiomyocytes.

### Neprilysin selectively alters SLN and PLB binding to SERCA

Since NEP protein was increased in a pattern that suggested SR localization, we considered how NEP activity might affect possible substrates in this organelle. A previous study suggested NEP cleavage of the SERCA regulator SLN reduced SLN-SERCA binding^8^, so we tested how NEP may alter PLB-SERCA binding as measured by fluorescence resonance energy transfer (FRET) between mCerulean3 labeled donor (mCer3-SERCA2a) and YFP labeled acceptor (YFP-PLB) in HEK293T cells. FRET was quantified on a cell-by-cell basis using automated fluorescence microscopy as previously described^19,36^. FRET between SERCA and PLB under control conditions was markedly reduced with NEP co-expression (**Figure 2A**), consistent with a decrease in PLB-SERCA binding. Since membrane protein-protein interactions may be sensitive to protein expression level, we compared each cell’s FRET efficiency with its YFP fluorescence intensity, taken as an index of PLB protein expression^37^. Plots of FRET efficiency vs PLB protein concentration showed that FRET between SERCA and PLB was lowest for cells expressing low levels of protein and increased to a maximal value (FRET_max_) in high-expressing cells, consistent with saturable binding of PLB to SERCA (**Figure 2B**). This relationship was well-described by a hyperbolic fit that yielded the apparent dissociation constant (K_D_), the protein concentration that yields half-maximal FRET efficiency (**Figure 2B**). Co-expression of NEP produced a qualitatively similar FRET binding curve, but we noted a distinct subpopulation of cells with markedly reduced FRET (**Figure 2C, red circle**). This lower-than-expected FRET efficiency suggests disruption of the PLB–SERCA interaction in these cells. Since the PLB-SERCA interaction seemed to be affected by coexpression of NEP, we examined the PLB sequence to determine possible cleavage sites. NEP is known to preferentially cleave at the C-termini of peptides with a preference for cleavage after valine residues^38^, so we examined whether PLB contains a candidate cleavage site. Sequence analysis revealed a conserved C-terminal valine (V49) in PLB and SLN (**Fig. S3, red box**), suggesting this residue may mediate NEP-dependent cleavage. To test the hypothesis that the valine residue at the C-termini of PLB was critical for NEP mediated cleavage, we mutated valine 49 of PLB to an alanine (V49A) and performed FRET analysis between SERCA (**Figure 2D**). In contrast to what we observed for WT-PLB, FRET from SERCA to V49A-PLB was unaffected by NEP coexpression, suggesting that V49 in PLB is critical for cleavage to occur (**Figure 2E**).

**Figure 2.**
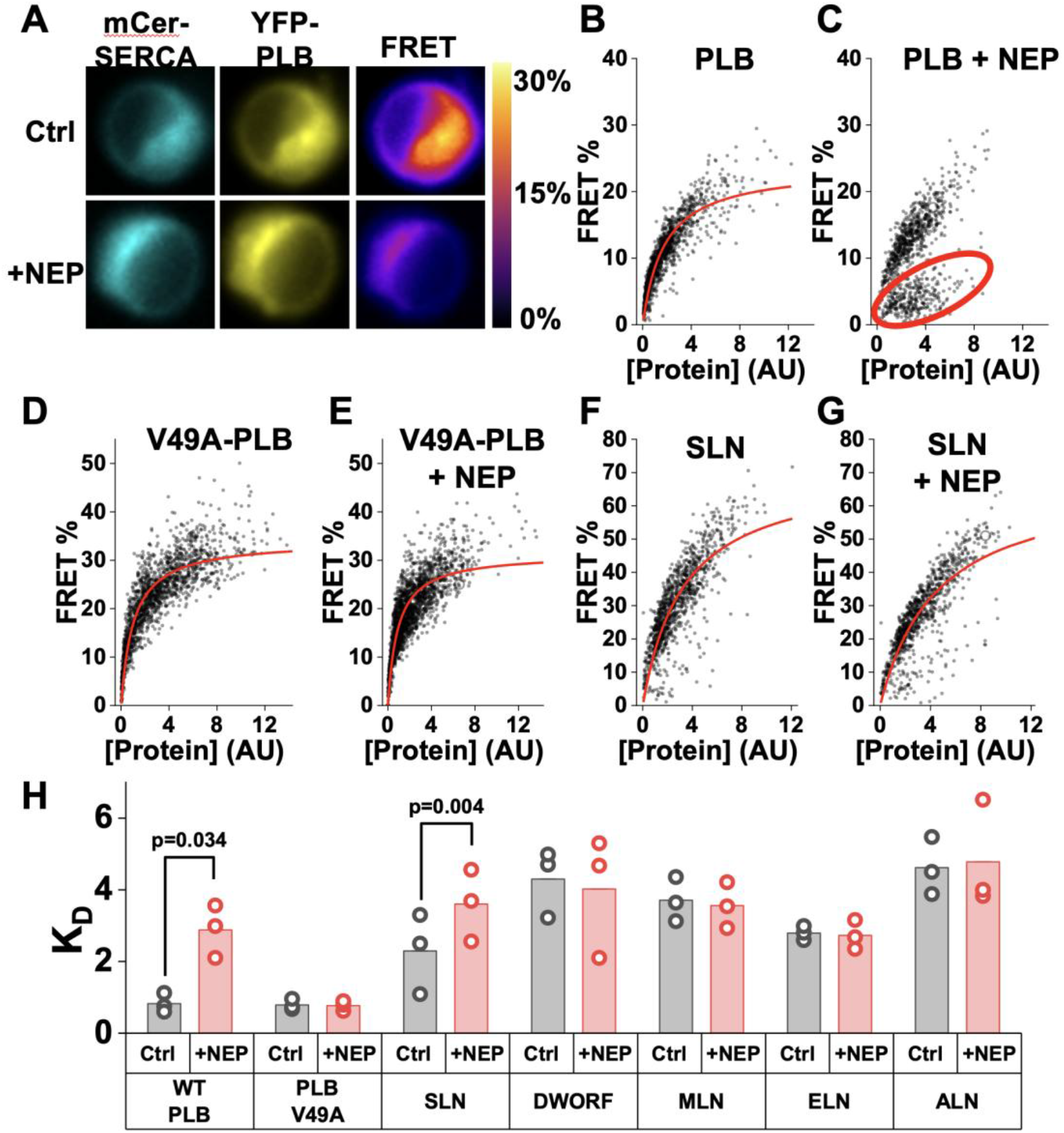
**A)** HEK293T cells transfected with mCer-SERCA, YFP-PLB (bottom row) NEP. **B)** Cell-by-cell FRET binding curve between mCer3-SERCA (donor) and YFP-PLB (acceptor). Apparent FRET efficiency is plotted as a function of acceptor expression. **C)** As in B, with NEP co-expression. The red oval highlights cells with markedly low FRET. **D)** FRET-based binding curve between mCer3-SERCA and V49A-YFP-PLB **E)** NEP co-expression did not affect FRET from SERCA to V49A-PLB. **F)** FRET binding curve between mCer3-SERCA and YFP-SLN, **G)** NEP co-expression reduced the apparent affinity of SLN for SERCA. **H)** Summary of apparent dissociation constants (K_D_) for all SERCA-binding micropeptides with and without NEP co-expression. Data represent 3 independent transfections, ∼1000 cells per transfection.

Previous studies have demonstrated that NEP can cleave the C-terminus of SLN, and this cleavage abolishes SLN inhibition of SERCA^8,39^. We quantified FRET between SERCA and SLN (**Figure 2F**), and found co-expression of NEP significantly increased the apparent K_D_ (**Figure 2G**), consistent with reduced binding affinity between the cleaved SLN product and the pump. Unlike the PLB–SERCA experiments, where NEP produced a distinct low-FRET subpopulation (**Figure 2C, red circle**), NEP shifted the entire SLN binding curve rightward, consistent with a uniform reduction in binding affinity between the cleaved SLN product and the pump. The PLB+NEP data are qualitatively different: rather than a uniform shift, a distinct subpopulation of cells exhibited markedly reduced FRET (**Figure 2C, red circle**), while the remaining cells tracked closely with control. Because the hyperbolic model does not capture this split population, the apparent K_D_ reported for PLB+NEP (**Figure 2H**) should be regarded as a qualitative index of disrupted binding rather than a true dissociation constant.

We further quantified whether NEP affected the binding affinity of other micropeptides in the regulin family known to modulate SERCA function^40^. Another-regulin (ALN), endoregulin (ELN), myoregulin (MLN) and dwarf open reading frame (DWORF) all showed avid binding to SERCA^41^, but the K_D_ measured for these proteins did not change with NEP co-expression. K_D_ values obtained for all micropeptides are summarized in **Figure 2H**; representative binding curves are provided in **Figure S4**. Comparison of the C-terminal sequences of these micropeptides (PLB, SLN, ALN, ELN, DWORF, and PLB), shows that only PLB and SLN contain a C-terminal valine residue at the predicted P1′ cleavage site (**Figure S3**), highlighting differences that may account for NEP substrate selectivity. Overall, the increased K_D_ of SLN and PLB and the emergence of a low-FRET subpopulation for PLB suggest a similar conclusion: NEP disrupts SERCA binding for both PLB and SLN, while leaving ALN, ELN, MLN, and DWORF unaffected. Increasing amounts of transfected NEP produced progressive increases in the apparent K_D_ of the PLB–SERCA interaction, indicating that increasing NEP levels directly weaken binding (**Figure S5A-E**).

### NEP-induced PLB cleavage and loss of membrane anchoring, is prevented by V49A-PLB mutation

Our previous work showed that removal of as few as three C-terminal residues disrupts PLB membrane anchoring and drives cytosolic mislocalization^24^. Because NEP cleaves the C terminus of SLN^8^, we hypothesized that the reduction in SERCA–PLB FRET after NEP coexpression may be due to loss of PLB membrane localization. To assess the relative partitioning of PLB in the ER membrane and cytoplasm, we selectively permeabilized the plasma membrane with saponin^42^ and monitored loss of YFP-PLB fluorescence from the permeabilized cell. PLB membrane retention was quantified by dividing residual YFP-PLB fluorescence (after saponin permeabilization) by the initial fluorescence before permeabilization. ER localization of YFP-PLB was unchanged by plasma membrane permeabilization (**Figure 3A**). As previously observed^24^, the truncation mutant YFP-I48X-PLB, diffused out of the cytosol into the surrounding medium following addition of saponin (**Figure 3A, bottom panel**). For YFP-WT-PLB cotransfected with NEP, we observed a significant loss of fluorescence after saponin permeabilization (**Figure 3A**), consistent with NEP-mediated cleavage and release of PLB from the membrane. In contrast, cells coexpressing NEP with V49A-PLB showed no loss of fluorescence after permeabilization (**Figure 3A**), indicating membrane retention. This observation underscores the importance of V49 for NEP recognition and cleavage of PLB. Quantification of these images is provided in **Figure 3B**. The data in **Figure 3A,B** were acquired by timelapse imaging of individual cells during saponin addition, which limited throughput. To assess whether NEP-mediated mislocalization was specific to PLB, we adopted a higher-throughput approach in which many fields of cells were imaged before and after saponin addition, quantifying the change in the fluorescence of each cell using an ImageJ macro (**Figure 3C**). This automated, high-throughput analysis confirmed that WT-PLB was solubilized by coexpression of NEP, whereas other micropeptides (including SLN) showed no detectable loss of intracellular fluorescence when coexpressed with NEP (**Figure 3C**).

**Figure 3.**
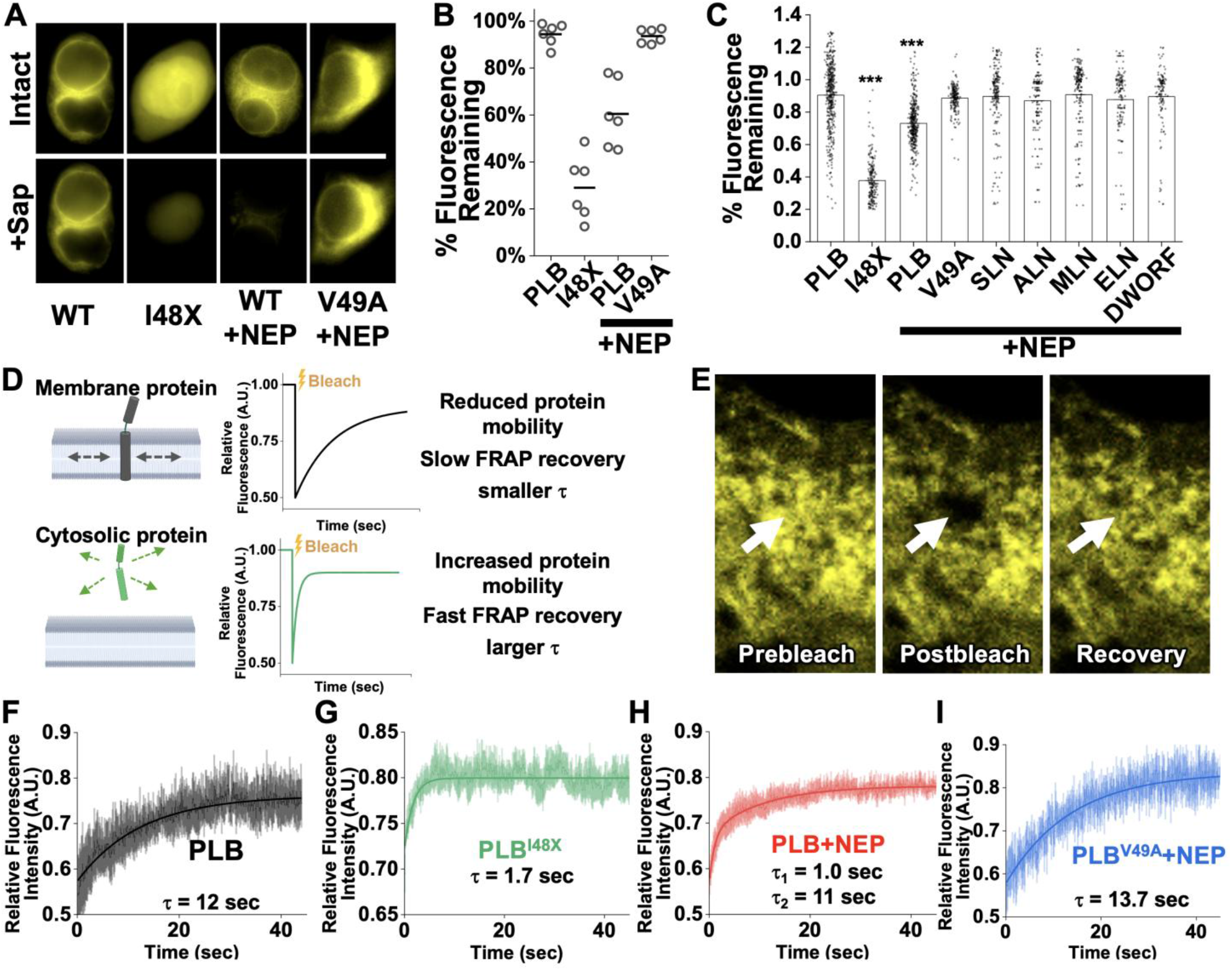
**A**) Fluorescence microscopy of cells before (intact) and after (+Sap) plasma membrane permeabilization with saponin revealed whether PLB was membrane anchored or soluble. **B)** Quantification of residual fluorescence after permeabilization (N/n = 3-6/7–9). **C)** Automated quantification of the loss of fluorescence of 150-250 cells after permeabilization. Only the truncated mutant I48X and PLB coexpressed with NEP were lost from permeabilized cells. Statistical significance was determined using one-way ANOVA followed by a Bonferroni post-hoc test. **D)** Schematic illustrating expected FRAP profiles of membrane-bound proteins versus freely-diffusible cytoplasmic proteins. **E)** TIRF image of YFP-PLB in ER membrane before and after spot photobleaching, and after 60 s of recovery. **F)** WT-PLB exhibited slow fluorescence recovery (τ ≈ 12 s), consistent with stable membrane association. **G)** The truncation mutant I48X-PLB showed rapid recovery (τ ≈ 1.7 s), reflecting increased lateral mobility. **H)** Coexpression of NEP with wild-type PLB produced biphasic recovery kinetics, with a fast component (τ_1_ ≈ 1.0 s) indicative of a mobile, cleaved species, and a slower component (τ_2_ ≈ 11 s) corresponding to intact PLB. **I)** NEP did not affect the fluorescence recovery of the cleavage-resistant mutant V49A-PLB (τ ≈ 8.8 s). FRAP traces are mean ± SD for 7-9 cells from 3 independent transfections.

Since NEP cleavage appears to release PLB from the membrane, we reasoned that cleaved PLB should exhibit increased intracellular mobility in intact cells. To quantify PLB diffusion we performed fluorescence recovery after photobleaching (FRAP) of YFP-PLB in live, intact cells. In a FRAP experiment, protein mobility is quantified by photobleaching fluorophores in a region of interest, then measuring the rate at which unbleached molecules diffuse into the bleached region and restore fluorescence. Faster fluorescence recovery indicates higher mobility (**Figure 3D**). Here, we used brief 514-nm laser illumination of a small spot to photobleach YFP-PLB. Representative images acquired before and after photobleaching and after 60 seconds of recovery are shown in **Figure 3E**. YFP-PLB exhibited slow fluorescence recovery (**Figure 3F**), consistent with slow diffusion of PLB in the membrane of the sarcoplasmic reticulum (SR). The recovery time (τ) corresponds to a diffusion coefficient (D) of 1.5 µm^2^/s, which is comparable to that of other membrane spanning micropeptides^43^. In contrast, the C-terminal truncation mutant YFP-PLB^I48X^ displayed rapid recovery (D = 12.3 µm^2^/s) (**Figure 3G**), indicating high mobility consistent with a freely diffusible cytosolic protein. Interestingly, coexpression of NEP with wild-type YFP-PLB resulted in a biphasic recovery profile that was best fit by a double-exponential function, revealing a fast component (D = 21.0 µm^2^/s) and a slower component (D = 1.0 µm^2^/s) (**Figure 3H**). The presence of this fast-diffusing fraction supports the hypothesis that NEP cleaves the C-terminus of PLB, generating a population of truncated, soluble products that diffuse freely in the cytoplasm. In contrast, fluorescent recovery of the cleavage-resistant mutant, V49A-PLB, displayed recovery kinetics resembling wild-type PLB (D ≈ 1.34 µm^2^/s) (**Figure 3I**), confirming mutation of V49 protects against NEP-mediated cleavage and maintains PLB membrane localization.

### Neprilysin enhances SR calcium uptake but reduces SERCA dynamic range

Since fluorescence microscopy revealed that NEP cleaves PLB and disrupts its binding to SERCA, we endeavored to determine how this may impact SERCA function. We assessed the functional consequences of NEP-mediated disruption of PLB using neonatal rat ventricular myocytes (NRVMs) as a model to assess SERCA-dependent Ca^2+^ reuptake. NRVMs were loaded with the ratiometric Ca^2^+ indicator Fura-2 AM, and the 340/380 nm fluorescence excitation ratio was recorded in cells expressing YFP-PLB alone (control) or coexpressing NEP and YFP-PLB (+NEP). Representative NRVMs expressing RFP α-actinin are shown in **Figure 4A**. Calcium transients from both conditions at 0.5 Hz pacing frequency are shown in **Figure 4B**. NEP-expressing NRVMs exhibited markedly increased calcium transient amplitudes relative to control cells (**Figure 4C**), along with a shorter time constant (τ) (**Figure 4D**). The markedly increased calcium transient along with a smaller τ were consistent with relief of PLB inhibition, and faster SERCA-mediated reuptake. We noted that relative τ, calculated as τ at 1 Hz divided by τ at 0.5 Hz, was significantly lower in NEP-expressing cells compared with control cells (**Figure 4E**), indicating a blunted force-frequency relationship (FFR), also known as treppe or the Bowditch Effect. Thus, cleavage of PLB by NEP blunts the physiological modulation of SERCA activity that normally adjusts calcium cycling in response to increasing pacing frequencies^44-46^. Taken together, these findings suggest that the functional consequences of NEP-induced cleavage of PLB are an acute increase in SR calcium load and contractility but loss of regulation of calcium handling. In particular, NEP cleavage of PLB compromises the Bowditch Effect that normally accelerates Ca^2^+ reuptake at higher heart rates.

**Figure 4.**
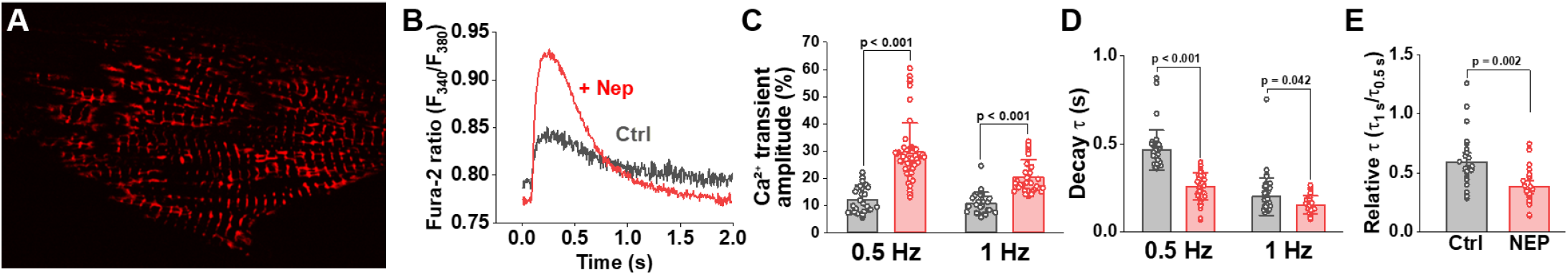
NEP enhances baseline Ca^2^+ cycling but reduces frequency-dependent SERCA modulation in NRVMs. **A)** NRVMs expressing RFP-α-actinin to mark sarcomere structures. Scale bar, 20 μm. **B)** Representative Ca^2^+ transients (F_340_/F_380_) at 0.5 Hz pacing. **C)** Ca^2^+ transient amplitude (ΔF_340_/F_380_) at 0.5 Hz. N/n = 8-14/3. **D)** Ca^2+^ transient decay time constant (τ) of cells paced at 0.5 Hz, fit by single-exponential decay. **E)** Ratio of decay τ at 1 Hz to decay τ at 0.5 Hz; larger values indicate increased frequency-dependent acceleration of Ca^2^+ reuptake. Differences were determined by two-way ANOVA with Tukey’s post hoc test (C, D) or unpaired t-test (E).

### Neprilysin Inhibition in Human iPSC-Cardiomyocytes

While the acute positively inotropic effect of NEP cleavage may help compensate for declining contractility in the failing heart, one would expect that the resulting loss of modulation of Ca^2+^ uptake would be deleterious *in vivo*. To determine if the net effect of NEP upregulation in DCM cardiac myocytes is adaptive or maladaptive, we performed transcriptomic analysis of human iPSC-derived cardiomyocytes (hiPSC-CMs) as a model of human myocardial gene expression^47,48^. The process of hypertrophy and failure was mimicked with endothelin-1 (ET-1) treatment, and the effect of NEP activity was evaluated by comparing transcriptome changes in the presence or absence of the NEP inhibitor sacubitrilat (Sac) (**Figure 5A**). An advantage of a cell culture model is that it isolates the direct, myocyte-specific effects of NEP inhibition from confounding effects of systemic changes such as circulating natriuretic peptides. RNA-sequencing of hiPSC-CMs revealed robust transcriptional changes in cells treated with ET-1 that resembled changes seen in human HF^49-51^ (**Figure 5B**), and we observed many differentially expressed genes in the comparison of cells treated with ET-1 versus NEP-inhibited cells treated with ET-1+Sac (**Figure 5B**). Gene ontology biological process (GO-BP) enrichment analysis classified the biological functions associated with the changes seen in **Figure 5B**. ET-1 treatment caused a downregulation of gene programs dominated by biosynthetic processes, including translation, ribosome biogenesis, and RNA metabolic processes (**Figure 5C, blue bars**), all frequently observed during remodeling in human HF^49,50,52^. ET-1 also triggered an upregulation of genes associated with canonical hypertrophic remodeling processes, including sarcomere organization, muscle contraction, and extracellular matrix–related organization (**Figure 5C, left, red bars**), which is also consistent with activation of a HF phenotype^49,50,52^. Furthermore, KEGG pathway analysis suggests that the genes change with ET1 treatment reflect cardiomyopathy (**Figure S6**). Interestingly comparison of ET-1 vs ET-1+Sac revealed upregulation in similar translational processes that were seen to be downregulated with ET-1 treatment, suggesting a protective effect of NEP inhibition. Biological processes that were significantly altered include regulation of calcium ion transport by the sarcoplasmic reticulum, actin filament depolymerization, cardiac muscle cell development, stress fiber assembly, and ATP metabolic process (**Figure 5C, right, blue bars**). Improvement of ATP metabolism further suggests NEP inhibition reduced the cellular stress response ^53^. The data suggest that NEP inhibition reduced markers of cellular stress by preventing PLB cleavage and restoring normal Ca^2+^ handling. **Figure 5D** summarizes changes in expression of genes considered to be hallmarks of human HF. Notably, ET1 increased *MME* (the gene which encodes neprilysin), suggesting that the disease-mimicking changes observed in the hiPSC-CM model is consistent with the NEP upregulation we observe in human DCM. ET-1 also increased expression of fetal gene markers (e.g., *NPPB* and *MYH7*) consistent with a hypertrophic response, and this change was ameliorated by Sac. Notably, while *PLN* mRNA increased with ET-1 and Sac restored a normal pattern, the other core calcium-handling transcripts (*ATP2A2, RYR2, CACNA1C*) were unchanged by either ET-1 or Sac. Thus, we hypothesize that NEP cleavage of PLB results in feedback activation of PLN transcription to compensate for NEP degradation of PLB protein, whereas inhibition of NEP by Sac rescues PLB protein levels and restores the normal PLB mRNA expression pattern. Together, these data establish that NEP inhibition reduces the hypertrophic response in cardiomyocytes, correcting changes in pathways linked to SR calcium transport, cytoskeletal remodeling, and metabolism.

**Figure 5.**
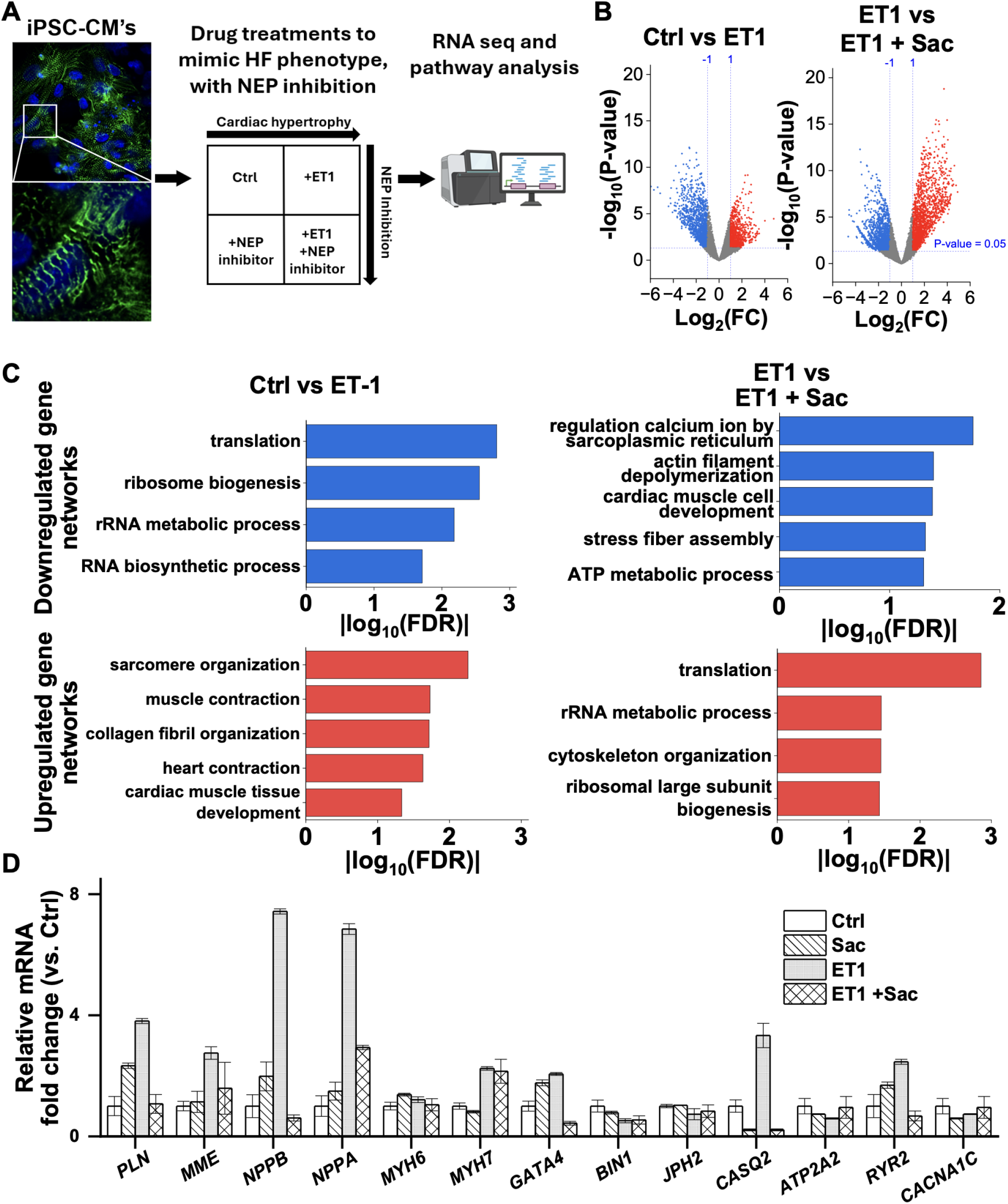
**A**) Summary of experimental design. **B)** Volcano plots showing differential gene expression changes for Ctrl vs ET-1 (left) and ET-1 vs ET-1 + Sac (right). ET-1 induces widespread transcriptional remodeling that was partially reversed by NEP inhibition. Significantly up- and downregulated genes (adjusted P < 0.05) are shown in red and blue, respectively. **C)** Gene ontology biological process (GO-BP) analysis. **D)** Relative expression of representative hypertrophic and Ca^2^+-handling genes. ET-1 induces fetal gene expression (e.g., increases in NPPA, NPPB, MYH7, decreases in JPH2, BIN1) and remodeling markers, whereas NEP inhibition partially normalizes these changes and preserves expression of Ca^2^+-handling components.

## Discussion

### A novel role for neprilysin in cardiac myocytes

The present work provides evidence for a novel pathogenic mechanism that may exacerbate declining cardiac function in heart failure. We propose that direct cleavage of PLB by NEP results in loss of PLB membrane anchoring, preventing it from regulating SERCA (**Figure 6A**). This mechanism may contribute to dysregulation of cardiac Ca^2^+ handling and loss of functional responsiveness to stress, both hallmarks of HF. **Figure 6B** summarizes a proposed cascade of consequences of PLB cleavage in the failing heart, beginning with pathogenic cardiac stress. This stress may arise from a variety of factors such as hypertension, genetic lesions, or metabolic dysfunction, any of which may initiate both adaptive and maladaptive responses in the heart^54^. In particular, upregulation of NEP in the SR and cleavage of PLB may represent a physiological response to stress that is initially compensatory. Relief of SERCA inhibition increases SR Ca^2+^ load and cardiac contractility (**Figure 6B-i**). However, this ostensibly adaptive response comes at the expense of dynamic range. PLB is a principal target of the β-adrenergic signaling cascade^55^, so excessive NEP activity is expected to reduce the heart’s responsiveness to neurohumoral control^56,57^ (**Figure 6B-ii**). Furthermore, PLB is critical for the Bowditch Effect, the intrinsic physiological mechanism that provides increased contractile force at faster heart rates^58^. Without PLB, the positive force-frequency response (FFR) is lost^46,59^ (**Figure 6B-iii**). Blunted β-adrenergic signaling and loss of FFR are defining characteristics of decompensated HF, and together they cause reduced contractile reserve (**Figure 6B-iv**). This functional decline imposes further cardiac stress, closing the maladaptive feedback loop that drives progression of disease^60^.

**Figure 6.**
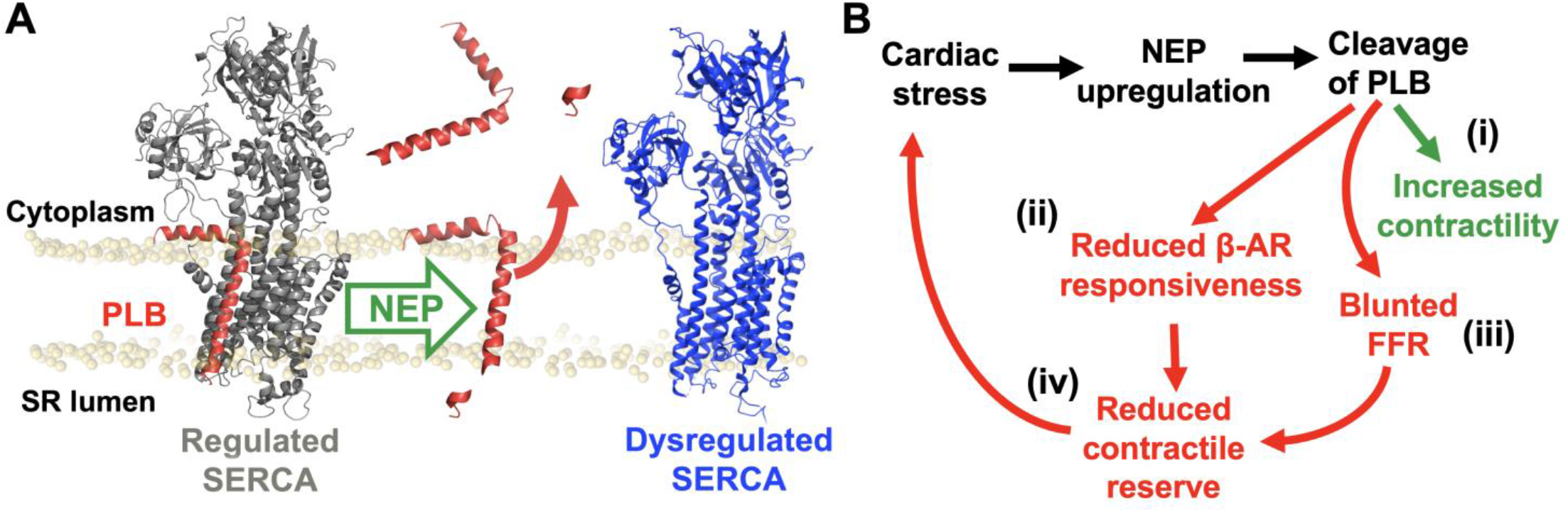
Proposed model for NEP-mediated dysregulation of SERCA in HF. **A)** PLB (red) anchored in the SR bilayer inhibits SERCA (gray); NEP cleaves PLB at its C-terminal valine, releasing the cytosolic fragment and yielding a dysregulated SERCA (blue) no longer subject to PLB-mediated control. **B)** A model of intracellular NEP activity in HF: cardiac stress drives NEP upregulation in cardiomyocytes and cleavage of PLB at the SR, acutely disinhibiting SERCA and increasing contractility (i, green) while progressively depleting the functional PLB pool, blunting β-adrenergic responsiveness (ii) and the force–frequency response (FFR) (iii) and reducing contractile reserve (iv), reinforcing cardiac stress in a maladaptive feedback loop.

### Targeting of membrane micropeptides by NEP

Schiemann et al. previously demonstrated that *Drosophila* neprilysin 4 cleaves the SERCA-inhibitory peptide sarcolamban at its luminal domain and showed that human NEP similarly hydrolyzes the luminal tail of SLN^8^. This proteolytic activity was proposed to serve a homeostatic role, supporting beneficial sarcolamban turnover and limiting SERCA inhibition in *Drosophila*. Our present findings suggest this mechanism is conserved in humans, where it acts on PLB, the predominant SERCA regulator in ventricular cardiomyocytes. We hypothesize that cleavage of PLB occurs at V49 based on the known recognition preference of NEP and the observation that the V49A mutation abolishes NEP-dependent disruption of PLB (**Figure 2, 3**). Of the SERCA-regulating micropeptides we tested, only PLB and SLN carry this C-terminal motif and only these two are substrates for NEP (**Figure S3**). Interestingly, SLN displayed complete retention within the ER even when coexpressed with NEP (**Figure 3**). We attribute this to SLN’s longer C-terminal tail that preserves its retention in the membrane even after excision of several C-terminal residues by NEP. However, we observed reduced binding of SLN to SERCA in cells coexpressing NEP. This is consistent with prior studies that showed the C-terminus of SLN is an important part of its SERCA regulatory function^61^. Thus, while NEP cleavage does not solubilize SLN as it does PLB, NEP can still modulate the SLN-SERCA regulatory interaction^61^.

### NEP inhibition ameliorates the hiPSC-CM hypertrophic response

Considering the cycle depicted in **Figure 6B** as a possible driver of cardiomyocyte decompensation, we reasoned that pharmacological NEP inhibition may interrupt the pathogenic feedback loop and slow the progression of disease. Indeed, application of the NEP inhibitor sacubitrilat (Sac) reduced upregulation of hypertrophic gene networks in a cell model of adverse cardiac remodeling, ET-1-treated hiPSC-CMs (**Figure 5B**). The reaction of these cells to ET-1 closely mirrors gene-expression changes observed in human HF^49,50,52^. NEP inhibition ameliorated this response and enriched salutary pathways related to Ca^2^+ handling, cytoskeletal organization, and metabolic processes. These results suggest that preventing NEP cleavage of PLB confers a cardioprotective phenotype in HF (**Figure 6B**). This framework also accounts for the rise in PLN transcript we observe in ET-1–stressed cells: NEP cleaves PLB protein, and the cell raises PLN transcription, possibly to compensate for the loss. When NEP is inhibited, PLB protein is preserved, the compensatory signal subsides, and PLN transcript normalizes. Importantly, because these experiments were performed in cultured cardiomyocytes lacking the systemic natriuretic peptide axis, this protective effect is intrinsic to cardiomyocytes.

### Implications for cardiac physiology and NEP inhibition

The present evidence for increased NEP activity in DCM accounts for the observation from our lab^30^ and others^27-29,62^ that PLB protein levels are reduced in failing myocardium. NEP-mediated cleavage of PLB at V49, releasing a soluble PLB from the SR membrane, provides a direct proteolytic mechanism for this reduction, as cleaved PLB in the cytoplasm is probably rapidly degraded. Similarly, a naturally occurring truncated PLB variant (L39X) was characterized by very low protein levels in heterologous expression and *in vivo*^*23*^. Moreover, we recently reported that disrupted proteostasis in HF generates toxic transmembrane peptides that compete with PLB for SERCA binding^30^, further eroding regulatory control. Together, NEP-mediated cleavage of PLB and competition from these aberrant peptides may contribute to decreased SERCA activity and loss of SERCA regulation characteristic of HF^63-65^. These findings suggest a novel mechanism of action for ARNI (sacubitril/valsartan) therapy. We propose ARNI therapy acts through a two-pronged mechanism to reduce cardiovascular death and HF hospitalizations in patients with chronic heart failure^66,67^: the established systemic action of sparing circulating natriuretic peptides, and a direct preservation ARNIs protect cardiac myocyte Ca^2^+ handling, that maintains the heart’s ability to dynamically respond to physiological challenge. Although the present study focused on specimens from patients with DCM, NEP mRNA is also elevated in hypertrophic cardiomyopathy^11^, suggesting ARNI therapy may offer a similar cardioprotective effect in other disease contexts. Future studies may provide additional rationale for expanding indications for ARNI therapy.

In summary, this study provides new mechanistic insight into the cardioprotective effects of NEP inhibition by identifying PLB as an intracellular substrate of NEP, revealing how NEP upregulation results in SERCA dysregulation in the failing heart.

## Supporting information

Major Resources Table

## Acknowledgments

The authors thank Prof. Aleksey Zima, Prof. Jonathan Kirk, [University of Chicago], Seby Edassery, and Dr. Elisa Bovo for technical assistance and helpful suggestions. The plasmid encoding neprilysin was generously provided by Prof. Heiko Meyer, [Universität Osnabrück].

## Materials and Methods

Details of the analyses and reagents, experimental models and subject details, experimental procedures, mass spectrometry–based proteomic analysis of human myocardial membrane fractions, fluorescence resonance energy transfer (FRET)-based binding assays, total internal reflection fluorescence (TIRF) microscopy and fluorescence recovery after photobleaching (FRAP), neonatal rat ventricular myocyte isolation and calcium imaging, human iPSC-derived cardiomyocyte culture and pharmacological treatment, bulk RNA sequencing and gene ontology analysis, and statistical analysis are included in the Supplemental Material.

## CRediT authorship contribution statement

**Jacob D. Cunningham:** Writing – Review & Editing, Writing – Original Draft, Visualization, Methodology, Investigation, Formal Analysis, Data Collection, Conceptualization. **Taylor A. Phillips:** Data Collection, Writing – Review & Editing. **Alexis R. Mazzenga:** Data Collection. **Kush N. Nagrani:** Data Collection. **Tuan H. Bui:** Data Collection. Seby Edassery: Data Collection and Data Analysis. **David Y. Barefield:** Conceptualization, Writing – Review & Editing. **Seth L. Robia:** Writing – Review & Editing, Supervision, Resources, Project Administration, Funding Acquisition, Formal Analysis, Conceptualization.

