## Supplementary Materials and Methods for "Neprilysin mediated cleavage of phospholamban dysregulates SERCA in heart failure"

This file includes:

Expanded Materials and Methods

Supplemental Tables S1

Supplemental Figures S1–S7

Supplemental References

### Materials and Methods

#### Immunofluorescence microscopy

Glass chamber slides (Nunc Lab Tek) were coated with poly-d-lysine (Millipore Sigma) and left to incubate at room temperature for 30 min. The poly-d-lysine was then aspirated, and the slides were left to air dry.

Approximately 25 mg of frozen human LV tissue was placed in a glass vial containing isolation solution (2 mM EGTA, 8.9 mM KOH, 10 mM imidazole, 7.1 mM $MgCl_2$, 5.8 mM ATP, 108 mM KCl) supplemented with 0.3% final volume Triton X-100 and protease inhibitor cocktails (Fisher Scientific). The tissue was homogenized 3 times 1 sec at 7,000 RPM with a mechanical homogenizer, passed through a 70-μm filter, and left to incubate on ice. After 20 min, the filtered homogenate was centrifuged at 120 *g* to pellet the myocytes. The cell pellet was resuspended in isolation solution free of Triton and pipetted onto the now dry poly-d-lysine-coated chamber slide and incubated at room temperature for 1 h and slides were washed with PBS.

Myocytes were fixed with ice-cold methanol for 1 min, followed by cold 4% paraformaldehyde for 3 min. To further permeabilize the cells, 0.5% Triton was added to each chamber and incubated for 20 min at room temperature, followed by two 15-min incubations with 0.1% Triton. A 0.1 M glycine solution (pH 7.4) was used for antigen retrieval by incubating the slides for 30 min at room temperature. The slides were next washed with PBS and incubated in blocking solution [1:1 vol/vol PBS to BSA solution (% BSA, 1% gelatin, 1% Tween-20, 0.001% $NaN_3$)]. After 1 h, the primary antibodies (anti-CD10, Abcam ab256494, 1:100; anti-SERCA2, Abcam ab2817) were added in blocking solution and slides were incubated for 12–14 h at 4°C.
Following incubation with primary antibody, the slides were washed with PBS and incubated with blocking solution containing secondary antibody (Abcam, AlexaFluor 488 or 568, 1:1,000, and Phalloidin 647) for 50 min at room temperature. The slides were mounted with Vectashield containing DAPI (Vector Laboratories), coverslipped, sealed with nail polish, and imaged at ×63 magnification on a Zeiss LSM 880 microscope under constant laser intensity and photomultiplier gain settings.

All super resolution imaging was performed at RT on a Zeiss LSM 880 Airyscan with a 63×1.4 NA oil-immersion objective in the “Airyscan” acquisition mode. Images were acquired using excitation lasers with wavelengths of 488 nm and 561(10) nm, and 647 nm with bandpass emission filters of 510(10) nm, 595(10) nm and 650(LP) nm respectively. Diode or argon laser at 0.9% power with 1.23 μsec pixel dwell, a 2x line averaging was used for all images acquired. Images were processed in Zen Blue with "joint Deconvolution" processing using 10 iterations for the Neprilysin, SERCA, and actin channels.

#### Western blotting

10 ug of membrane enriched fractions used in mass spectrometry were diluted in 4× Laemmli sample buffer with β-mercaptoethanol at 1:1 ratio, denatured at 90 °C for 5 min, run on a 4-15% polyacrylamide gradient gel, and transferred to a polyvinylidene difluoride membrane. The membrane was stained with Revert total protein stain (LI-COR Biosciences) for 5 min to obtain total protein in each lane and then blocked for 1 h at room temperature in Intercept blocking buffer (LI-COR Biosciences) diluted at a 1:1 ratio in PBS with Tween 20 (PBS-T). Blots were incubated overnight at 4 °C with primary antibody rabbit anti-CD10 (human, ab256494, 1:1000, Abcam). The blots were then incubated with anti-rabbit (IRDye 800CW; LI-COR Biosciences) secondary antibody diluted 1:10,000 in PBS-T. Blots were imaged using an Azure c600 gel imaging system and analyzed using the ImageJ.

### Mass Spectrometry

*Sample preparation*

Left ventricle samples from non-failing and DCM human hearts were flash frozen and stored at -80°C, previously. Non-failing samples were taken from rejected donors with cause of death due to non-cardiovascular related issues and DCM samples were from end stage heart failure patients with EF<40% per NYHA guidelines. Samples were thawed and homogenized in SR prep buffer (100 mM KCl, 2.5 mM $K_2HPO_4$, 2.5 mM $KH_2PO_4$, 2 mM EDTA) containing protease inhibitors. Samples were centrifuged at 6,400 *g* for 20 minutes at 4°C. The supernatant containing soluble proteins from the previous step was centrifuged at 10,000 *g* for 20 min to remove heavy debris. Subsequently, the supernatant was centrifuged at 9,700 *g* for 20 min at 4. The supernatant was then collected, and further spun at 86,000 *g* at 4 °C for 60 min. Finally, the pellet containing the membrane proteins was dissolved in 100 µl of buffer B (1 M sucrose, 50 mM KCl) and stored at −80 °C. The total protein concentration was determined using Bicinchoninic Acid Kit for Protein Determination (Pierce), and all reagents were purchased from Sigma Aldrich if not stated otherwise.

Proteins from the membrane enrichment were extracted using the following protocol[32]. Briefly, 100 ug of five biological replicates samples of non-failing and DCM membrane fractions were incubated in 1 ml of chloroform on a shaker at room temperature for 1 h. To this, 1 ml of methanol-water (1:1 vol/vol) was added, and the mixture was vortexed vigorously for 30 min. The mixture was spun at 2,000 *g* for 1 min, and the chloroform layer was discarded. Another 1 ml of chloroform was added to further extract lipids. The mixture was sonicated in a bath sonicator for 30 min without heating. Ice-cold water was added at regular intervals to prevent heating due to continuous sonication. The mixture was then spun at 10,000 *g* for 5 min. To the aqueous layer and the insoluble interface, four times the volume of acetone was added and incubated at 4°C for 1 h. The protein was collected by pelleting at 10,000 *g* for 5 min. The protein pellet was washed twice with acetone, dried on ice, and dissolved in 2M urea and 250 mM ammonium bicarbonate to complete dissolution. Protein denaturation and reduction were carried out by adding DTT to 10 mM and incubated for 30 min at 50°C, and alkylation was carried out by the addition of iodoacetamide at 30 mM final concentration and incubated at 37°C for 1 h in the dark. Protein was then trypsin/LysC digested in 30% methanol at 20:1 (wt/wt) protein to protease ratio and incubated overnight at 37°C and dried using a speedvac.

### Proteomics

Purified peptides, 500 ng, were loaded onto a Vanquish Neo UHPLC system (Thermo Fisher) with a heated trap and elute workflow with a c18 PrepMap, 5mm, 5uM trap column(P/N 160454) in a forward-flush configuration connected to a 50 cm Easyspray analytical column(P/N ES75500PN) 2uM,100A,75um x 500mm ,with 100% Buffer A (0.1% Formic acid in water) with flow rate of 0.300 µL and the column oven operating at 35 °C. Peptides were eluted over a 150 min gradient, using 80% acetonitrile 0.1% formic acid (buffer B), going from 4 % to 10% over 10 min, to 35% over 100 min, then to 50% over 22 min, then to 99% over 6 min and kept at 99% for 12 min, after which all peptides were eluted. Spectra were acquired with an Orbitrap Eclipse Tribrid mass spectrometer (Eclipse) with FAIMS Pro interface (Thermo Fisher Scientific) running Tune 3.5 and Xcalibur 4.5 was used for all acquisition methods, spray voltage set to 1600V, and ion transfer tube temperature set at 300oC, FAIMS switched between CVs of −45 V, – 55 V, and −65 V with cycle times of 1.5 s. MS1 spectra were acquired at 120,000 resolutions with a scan range from 375 to 1600 m/z, normalized AGC target of 300%, and maximum injection time set to auto, S-lens RF level set to 30 without source fragmentation and datatype positive and profile; Precursors were filtered using monoisotopic peak determination set to peptide MIPS; included charge states, 2-7 (reject unassigned); dynamic exclusion enabled, with n = 1 for 60s exclusion duration at 10 ppm for high and low. DDMS2 scan using isolation mode Quadrupole, Isolation

Window (m/z): 1.6; Activation Type set to HCD with 30% Collision Energy (CE), Detector Type: Ion Trap; Scan Rate: Turbo; AGC Target: 10000; Maximum Injection Time:35 ms, Microscans: 1 and Data Type: Centroid.

**MS Data Analysis**
Raw data were analyzed using Proteome Discoverer 2.5 (Thermo Fisher) using Sequest HT search engines. The data were searched against the Human entries in the UniProt protein sequence database (Homo sapiens, Proteome ID UP000005640). The sequest search parameters included precursor mass tolerance of ten ppm and 0.6 Da for fragments, 2 missed trypsin cleavages, oxidation (Met) and acetylation (protein N-term) as variable modifications, and carbamidomethylation (Cys) as a static modification. Percolator PSM validation was used with the following parameters: strict false discover rate (FDR) of 0.01, relaxed FDR of 0.05, maximum ΔCn of 0.05, and validation based on q-value. Precursor abundance was quantified using intensity values. Data were normalized based on total peptide amount, with no additional scaling applied. Protein abundances were calculated by summing peptide intensities. Abundance ratios between groups were determined using a class comparison approach with a background-corrected t-test. Differentially expressed proteins were defined by a $p$-value < 0.05 and an absolute log2 fold change greater than 1.0.

**Homology alignment**
Amino acid sequences for human phospholamban (PLN, UniProt P26678), sarcolipin (SLN, O00631), myoregulin (MRLN, P0DMT0), DWORF (P0DN84), endoregulin (ERLN, P0DI80), and another-regulin (ALN, Q8WVX3) were obtained from the UniProt Knowledgebase (UniProtKB). All sequences were verified to correspond to canonical human isoforms. Sequences were imported into Jalview (v2.11.3.2) using the “Input Alignment → From Textbox” function and formatted in FASTA style. Multiple sequence alignment was performed via the integrated Clustal web service with default protein parameters (BLOSUM62 substitution matrix, gap opening = 10, gap extension = 0.2). Alignment outputs were visualized in Jalview’s ClustalX color scheme to highlight residue conservation, charge, and hydrophobicity. Residue-wise conservation, consensus, and buriedness scores were calculated in Jalview using the built-in “Conservation” and “Buried Index” tools. The buried index reflects amino-acid propensity for membrane-embedded positions.

**Plasmid constructs**
mCerulean3 (mCer3) or enhanced yellow fluorescent protein (YFP) was fused to the N terminus of each respective micropeptide or the human isoform of SERCA2a (GenBank™ Accession Number P16615) as previously described[42, 66, 67]. Genes encoding human isoforms of PLB, SLN, ELN, ALN, DWORF and MLN were fused to EYFP via a five or seven amino acid linker (of sequence SGLRS or GGGGGKL respectively) to the N terminus of the micropeptide as previously described[38]. The I48X deletion and valA substitution mutant of PLB was generated as previously described[16]. We previously have shown that PLB fused with a fluorescent tag does not alter its ability to regulate the SERCA pump[42].

The unlabeled human neprilysin construct was a generous gift from Heiko Meyer (Universität Osnabrück) and was generated as previously described[6]. For localization controls, mCherry was fused to either the N or C terminus of NEP. All functional experiments used the unlabeled NEP construct, which carries no fluorescent protein tag. Notably, in those heterologous expression experiments we determined that N-terminal tagging caused mislocalization of NEP to the cytoplasm of HEK293T cells (**Figure S2**). An alternative C-terminal mCherry fusion tag preserved NEP localization but abolished its enzymatic activity in subsequent functional assessments. Accordingly, subsequent functional experiments were performed with NEP with no fluorescent protein tag.

**Cell Culture and Transfection**

HEK293T cells (Agilent, Santa Clara, CA, USA) were cultured in Dulbecco's modified Eagle's medium (DMEM) cell culture medium supplemented with 10% fetal bovine serum (ThermoScientific). Following culture at 20-30% confluency, the cells were transiently transfected using Lipofectamine 3000 transfection kit (Invitrogen) as per instructions provided. Forty-eight hours post transfection, cells were trypsinized (Thermo Fisher Scientific, Waltham, MA, USA) and replated onto poly-D-lysine– coated 2-well glass bottom chamber plates and allowed to settle down for 1 hour before imaging, or 12 hours prior to imaging if cells were permeabilized in the experiment. Prior to imaging, cells were incubated at 37°C in DMEM with 10% fetal bovine serum. Our previous studies of DWORF and PLB suggest that the expression levels achieved in AAV-293 cells are somewhat lower than native myocardium[42].

**Fluorescence Resonance Energy Transfer (FRET) Quantification**

Acceptor sensitization FRET was quantified as previously described[34]. Briefly, HEK293T cells were transiently transfected with mCer3-donor and eYFP acceptor–labeled Cells were transfected with FRET-binding partners in a 1:5 M plasmid ratio, with or without 2.5 ug of unlabeled NEP, and seeded on poly-D-lysine coated chamber slide 1 hour prior to imaging. FRET acceptor sensitization was measured by automated fluorescence microscopy using an inverted microscope (Nikon Ti Eclipse 2). A Lumencor Spectra X excitation system was used to excite samples with 50 ms exposure time. Emitted light was passed through a triple band dichroic filter (CFP/YFP/mCherry Spectra X emission filter set) before detection with a Photometrics Prime 95B camera. 3 image channels were obtained for each field: mCer3 donor (excitation: 430(24) nm, detection: 475/25 nm), YFP acceptor (excitation: 500(20) nm, detection: 535(30) nm), and FRET (excitation: 430(24) nm, detection: 535(30) nm). 144 images (~1000 total cells per condition) were collected from three independent experiments with a 40 × 0.75 numerical aperture objective: mCer3, YFP, and FRET (Cer excitation/YFP emission) using the Elements software. Cells expressing mCer3 had an area of 136 to 679 $\mu m^2$, and were at least 40% circular were automatically scored for Cer, YFP, and FRET fluorescence intensity with a rolling background subtraction using a plugin in Fiji (imagej.net/software/fiji/). Fluorescence intensities from mCer, YFP, and FRET channels ($I_{DD}$, $I_{AA}$, and $I_{DA}$ respectively) were used to calculate sensitized emission FRET according to $E_{app} = F_c/(F_c + G \times I_{DD})$, where $F_c = I_{DA} - (a \times I_{AA}) - (d \times I_{DD})$, where $E_{app}$ represents the apparent FRET efficiency corrected for imaging-induced photobleaching, and $F_c$ represents the sensitized emission FRET intensity corrected for crosstalk between channels. Parameters $a$ and $d$ are crosstalk constants calculated as $a = I_{DA}/I_{DD}$ for a control sample transfected only with the YFP acceptor and $d = I_{DA}/I_{AA}$ for a control sample transfected only with the mCer donor. $G$ is the ratio of sensitized acceptor emission to a corresponding amount of donor recovery in the $I_{DD}$ channel after acceptor photobleaching ($I_{DD}^{post}$), defined by the equation $G = F_c/(I_{DD}^{post} - I_{DD})$. For the experiments in this study using mCer3 and YFP FRET pairs, these values were determined to be $a = 0.1853$, $d = 0.4051$, and $G = 2.78$. Apparent FRET efficiencies for each cell were then plotted as a function of YFP acceptor fluorescence intensity, which was used as an index of relative protein expression.

**Quantification of loss of fluorescence after saponin addition**

To quantify the solubilization of PLB, cells were cotransfected with Cer-SERCA and YFP-PLB truncation mutants at a 1:1 ratio and subjected to fluorescence imaging using an inverted Nikon Ti2E microscope with a 40x (for automated analysis) or 80x objective. mCer3 and YFP were sequentially excited at 458 and 514 nm, respectively. Cells were then permeabilized with 100 μg/ml saponin in 120 mM, KCl 15 mM, $KH_2PO_4$ 5 mM, $MgCl_2$ 0.75 mM, dextran 2%, ATP 5 mM, HEPES 20 mM, and EGTA 2 mM, pH 7.2, and the same fields were reimaged after permeabilization. Loss of YFP signal following saponin treatment was used as a measure of PLB solubilization. The residual YFP/Cer signal was quantified per cell, and solubilization was expressed as the fractional decrease in YFP signal relative to the pre-permeabilization image. Automated analysis at 40× was performed on ~1,000 cells across three independent transfections; higher-resolution 80× imaging was performed on 8 cells across three independent transfections.

**Total Internal Reflection (TIRF) Microscopy and Fluorescence Recovery after Photobleaching (FRAP)**

Total internal reflection fluorescence (TIRF) was performed as previously described[40]. Briefly, imaging was performed using the 457.9 nm line of laser, coupled through the objective via a multiband dichroic mirror. Emission under TIRF illumination was collected using the filter sets described above. For spatially localized, acceptor-specific photobleaching experiments, the laser line was isolated with a line-selective filter and delivered to the sample through a 10/90 beam splitter. The photobleaching beam was spatially confined by focusing it onto the specimen using a Keplerian telescope consisting of two planoconvex lenses. Photobleaching was achieved with a brief laser exposure of approximately 500 ms. Post-bleach images were averaged and normalized to the corresponding pre-bleach image to generate $F/F_0$ time series. The spatial extent of the bleached region in the acceptor channel and the corresponding donor dequenching were quantified using MetaMorph software by measuring fluorescence intensity along a line transecting the region of interest.

**NRVM Isolation, Transfection, and $Ca^{2+}$ Imaging**

All animals were housed and sacrificed in accordance with Loyola University Chicago's Institutional Animal Care and Use Committee in adherence to the US National Institutes of Health Guide for Care and Use of Laboratory Animals. Neonatal rat ventricular myocytes were isolated from 0 to 1-day old Sprague-Dawley pups (Charles River Laboratories) as described[68]. Briefly, ventricles were dissected, minced, and digested at 37 °C in Krebs–Henseleit buffer containing collagenase and 0.05% trypsin for six sequential rounds. At each step, the digest was collected, filtered, and quenched in DMEM supplemented with 10% FBS and 1% penicillin–streptomycin. Pooled cells were centrifuged at 800 × g for 10 min, resuspended in isolation media, and pre-plated for 1.5 h at 37 °C to remove fibroblasts. The non-adherent cardiomyocyte-enriched suspension was collected, centrifuged at 500 × g for 10 min, and resuspended in maintenance media (DMEM/F12, 18.5% M199, 5% horse serum, 1% FBS, 1% penicillin–streptomycin, 1% insulin-transferrin-selenium, 10 μM BrdU). Final suspensions were adjusted to $1 \times 10^6$ cells/mL for plating on coverslips.

NRVMs were then transfected using the Lipofectamine 3000 transfection kit (Invitrogen) using 0.5 ug of YFP-PLB and 1.5 ug of NEP. Twenty-four hours after transfection, cells were rinsed with Tyrode's buffer and incubated for 13 – 15 minutes in Tyrode's buffer with 1 μM Fura-2 AM (ThermoFisher) at room temperature. After incubation, coverslips were washed with Tyrode's free of Fura-2 AM for an additional 15 minutes at RT and then transferred to a recording chamber.

GFP positive cells were first located, and all recordings were performed under electrical stimulation at 0.5 and 1 Hz for 5 ms and 30 V (IonOptix MyoPacer EP—Field Stimulator). The range of fluorescence excitation was 340/380 nm and collected by a photomultiplier tube via the

40x objective at 510 nm with a 75 W Xenon lamp. The IonOptix contractility system was used to quantify calcium transients and data were processed using IonWizard. Calcium transients were averaged and plotted using OriginPro (Version 2025, OriginLab Corporation, Northampton, MA, USA).

**Differentiation and Culture of hiPSC-Derived Cardiomyocytes**

Human WTC induced pluripotent stem cells carrying a monoallelic, C-terminal mEGFP tag at the endogenous ACTN2 locus (AICS-0075-085, clone 85; Allen Cell Collection, obtained through the Coriell Institute for Medical Research) were differentiated into cardiomyocytes using temporal modulation of Wnt signaling. hiPSCs were initially seeded onto a Matrigel-coated (Corning) plate in a solution consisting of mTeSR™ Plus culturing media (STEMCELL Technologies) and 0.1% thiazovivin (STEMCELL Technologies). The following day, the cells would be refed and maintained with just mTeSR™ Plus until they reached around 70%-80% confluency (Day 0), where then they were cultured in a medium consisting of RPMI 1640 (Gibco), B-27™ Supplement minus insulin (Gibco), and 7.5 µM CHIR 99021 (Tocris Bioscience). On Day 2, the medium was replaced with RPMI 1640, B-27™ Supplement minus insulin, and 5 µM of IWP 2 (Tocris Bioscience). On Day 4, the medium was changed to just RPMI 1640 with just B-27™ Supplement minus insulin. For Days 6-14, the cells would be fed with RPMI 1640 with regular B-27™ Supplement (Gibco) (RPMI+B27) every other day until spontaneous contractions start to occur. Cardiomyocyte selection would then be initiated by changing the medium to a solution consisting of RPMI 1640, no glucose (11879020, Gibco), B-27™ Supplement minus insulin, and 10 mM sodium L-lactate (L1450014, Thermo Scientific Chemicals). After 2 days post-selection, the iPSC-cardiomyocytes were then disassociated with 10X TrypLE (Gibco) for 5-10 minutes at 37 °C and then centrifuged at 1000 × g for 5 minutes. The supernatant was then aspirated, and the cell pellet resuspended in a solution consisting of RPMI+B27, 1% penicillin–streptomycin (Gibco), 0.1% thiazovivin, and 10% KnockOut™ Serum Replacement (Gibco) before then being seeded onto a Matrigel-coated plate at a concentration of $3 \times 10^6$ cells/mL. The following day, the cardiomyocytes would be refed and maintained with just RPMI+B27.

**RNA-Sequencing**

Bulk 3′ RNA sequencing was performed by Plasmidsaurus using a stranded, single-end workflow on an Illumina NovaSeq X Plus instrument. Further detailed gene set enrichment and functional association analyses were performed using the GO Functionome web resource[69], which integrates protein–protein interaction networks with Gene Ontology annotations.

**Statistics and reproducibility**

All experiments were performed with independent biological replicates as indicated in the figure legends and/or below. Human left ventricular (LV) tissue was obtained from non-failing (NF) donors and dilated cardiomyopathy (DCM) patients as described in Methods. For membrane-enrichment proteomics, five biological replicates per group were analyzed (NF, n=5; DCM, n=5). Western blot validation used membrane-enriched fractions from the same cohort (NF n=5, DCM n=5). For cell-based assays (HEK293T, NRVMs, hiPSC-CMs), each experiment was repeated on ≥3 independent days unless otherwise indicated, using fresh transfections/cultures each day. "n" denotes independent samples (hearts, culture preparations, or experimental days) as specified; when single-cell imaging was used, the number of analyzed cells per condition is reported and nested within biological replicates.

**Software**

Image analysis was performed in Fiji/ImageJ (including custom macros where stated). Curve fitting and statistics were performed in OriginPro. Proteomics was analyzed in Proteome

| | Non-Failing | Heart Failure |
|---|---|---|
| Age | 52.3±8.1 | 43.3±10.4 |
| Male | 3 (60) | 4 (80) |
| Female | 2 (40) | 1 (20) |
| African American, n (%) | 1(20) | 2 (40) |
| NT-proBNP — ng/liter [pg/mL, median (IQR)] | NA | 2090.5 [1708-2539] |
| EF (%) ± SD | NA | 13 ± 8 |

**Supplementary Table 1.** Clinical and demographic characteristics of human left ventricular tissue donors. Summary of donor characteristics for non-failing (NF, n=5) and end-stage heart failure (HF, n=5) left ventricular tissue specimens used for proteomic and western blot analyses (Figure 1). Continuous variables are presented as mean ± SD, except NT-proBNP, which is reported as median with interquartile range. Categorical variables are reported as n (%). Non-failing samples were obtained from donors whose hearts were rejected for transplantation due to non-cardiovascular causes of death; heart failure samples were obtained from end-stage HF patients with ejection fraction <40% per NYHA criteria. NT-proBNP and ejection fraction (EF) were not available (NA) for non-failing donors. IQR, interquartile range; NT-proBNP, N-terminal pro-B-type natriuretic peptide; EF, ejection fraction.

Discoverer v2.5 with Percolator validation as described in Methods above.

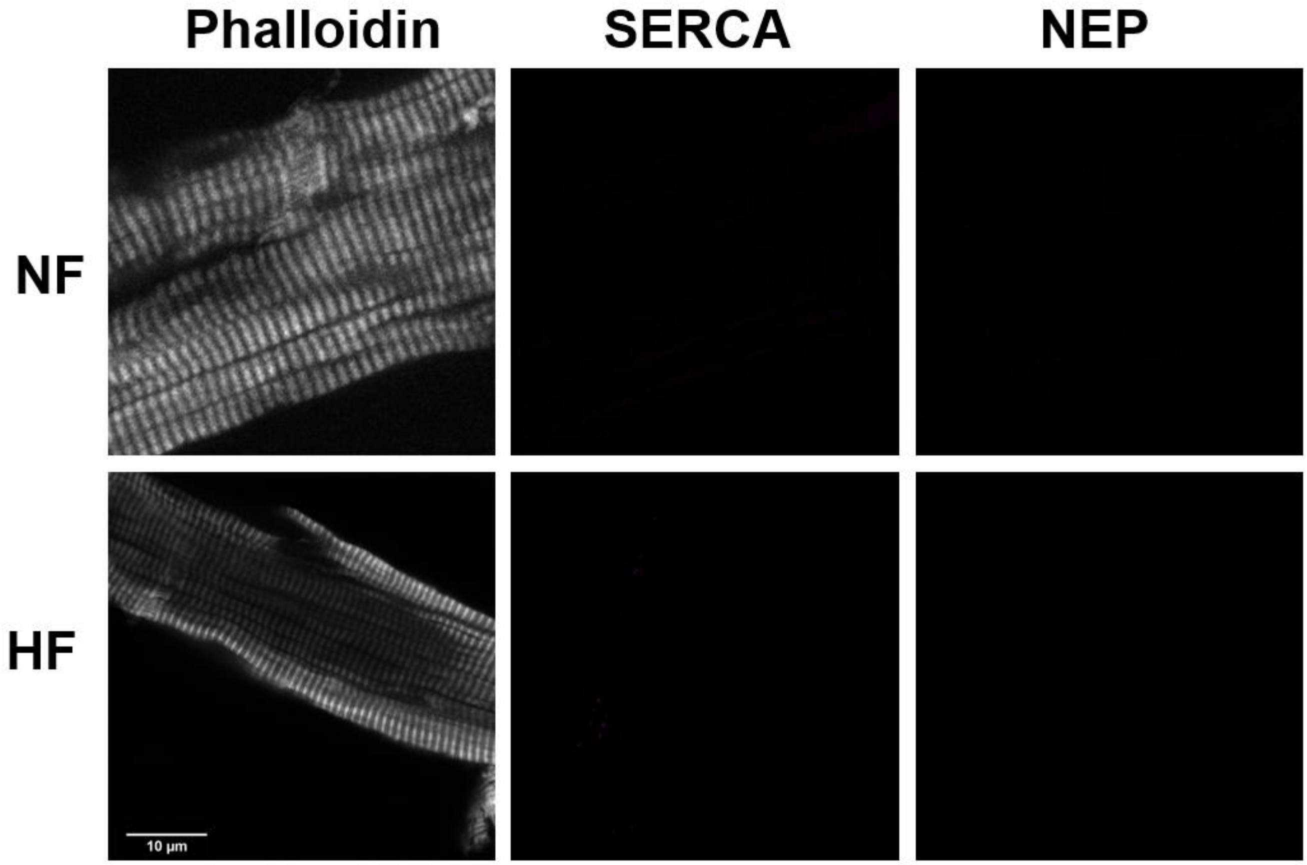


**Figure S1. Secondary antibody–only control for NEP immunofluorescence in human myocardium.** Representative confocal images of non-failing (NF) and heart failure (HF) human left ventricular cardiomyocyte preparations processed identically to Figure 1F but with primary antibodies omitted. AlexaFluor 488 and AlexaFluor 568 secondary antibodies, phalloidin, and DAPI were applied as described in Methods. Absence of punctate signal in both NF and HF samples confirms that the NEP and SERCA puncta in Figure 1F reflect specific antibody binding

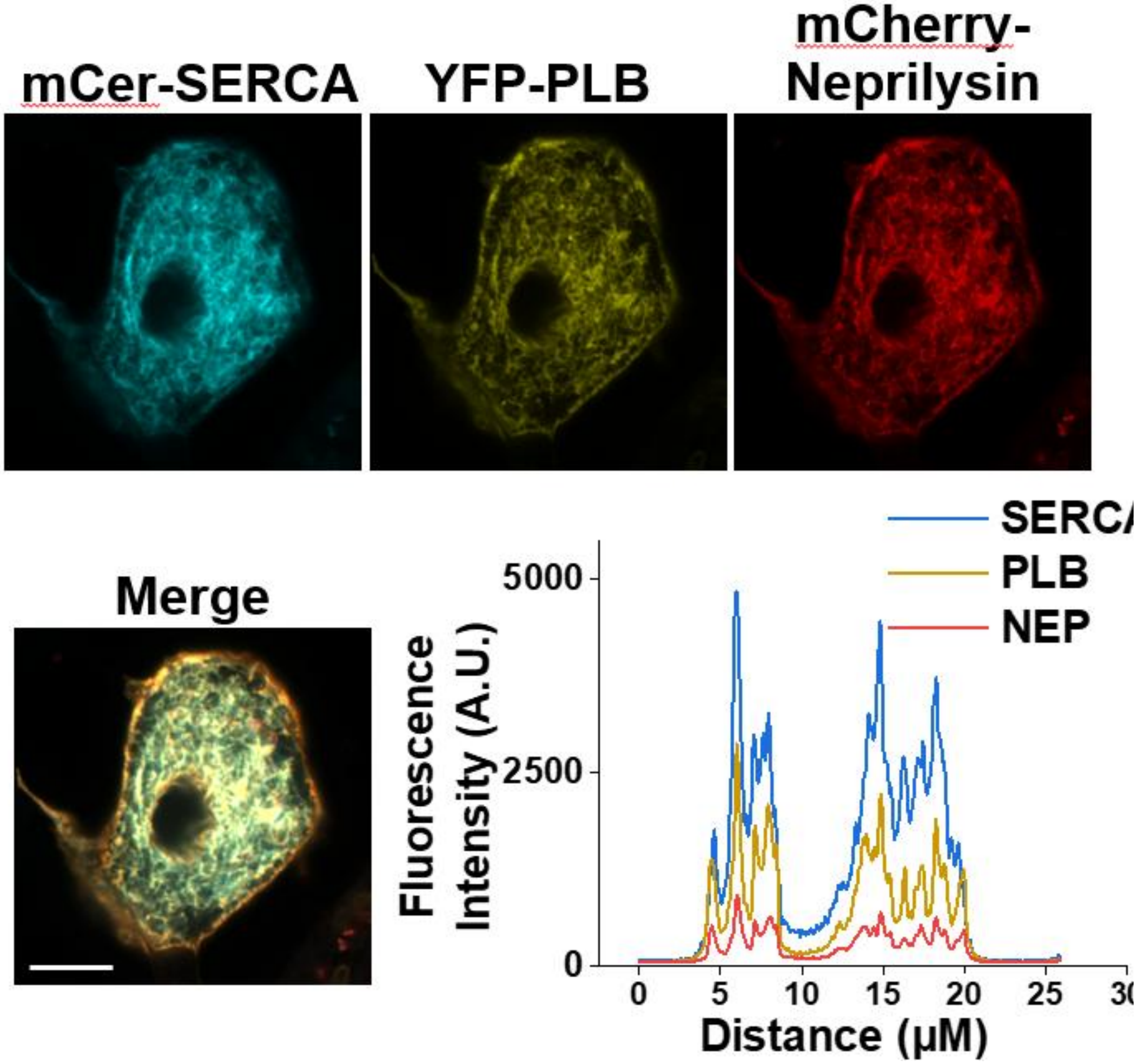


**Figure S2. Heterologous expression and localization of tagged NEP constructs in HEK293T cells.** **(A)** Representative confocal images of HEK293T cells co-expressing untagged NEP, mCer3-SERCA, and YFP-PLB. NEP was visualized by anti-CD10 immunostaining. NEP signal colocalizes with both SERCA and PLB at the endoplasmic reticulum, consistent with intracellular SR-like localization observed in cardiomyocytes (Figure 1F). **(B)** N-terminal mCherry-NEP fusion mislocalizes to the cytoplasm rather than membrane compartments. **(C)** C-terminal NEP-mCherry fusion preserves membrane localization but abolishes enzymatic activity in [substrate cleavage / SERCA-PLB FRET

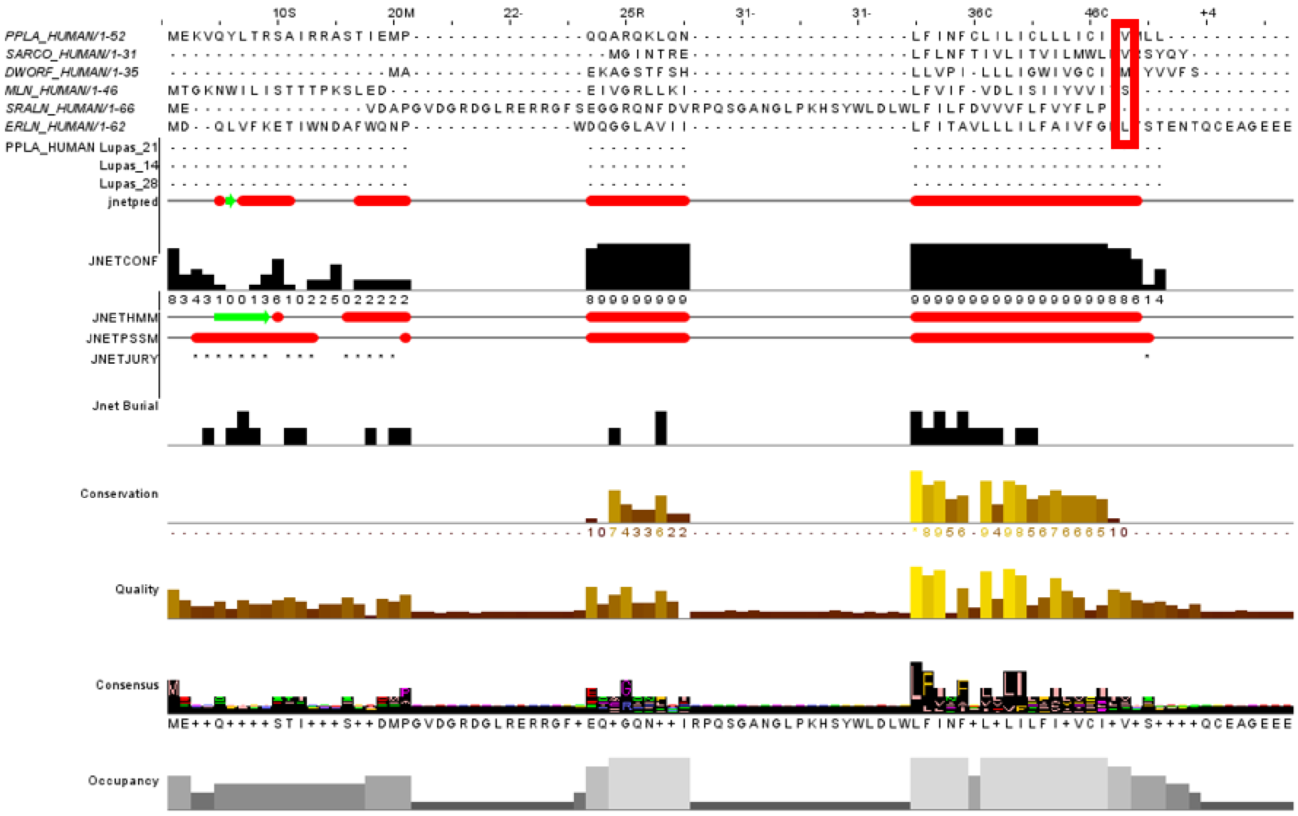


**Supplementary Figure 3.** Multiple sequence alignment of SERCA-regulatory micropeptide C-termini. Alignment of human phospholamban (PLN/PLB, UniProt P26678), sarcolipin (SLN, O00631), myoregulin (MRLN, P0DMT0), another-regulin (ALN, Q8WVX3), endoregulin (ERLN, P0DI80), and DWORF (P0DN84), generated in Jalview using Clustal with default parameters (BLOSUM62, gap open 10, gap extend 0.2). The conserved C-terminal valine present in PLB (V49) and SLN — and absent in MLN, ALN, ELN, and DWORF —highlighted by the red box. Coloring reflects the ClustalX scheme; conservation, consensus, and buried-index tracks shown below the alignment. The P1′ valine identifies PLB and SLN as the only members of this family with the canonical NEP cleavage motif.

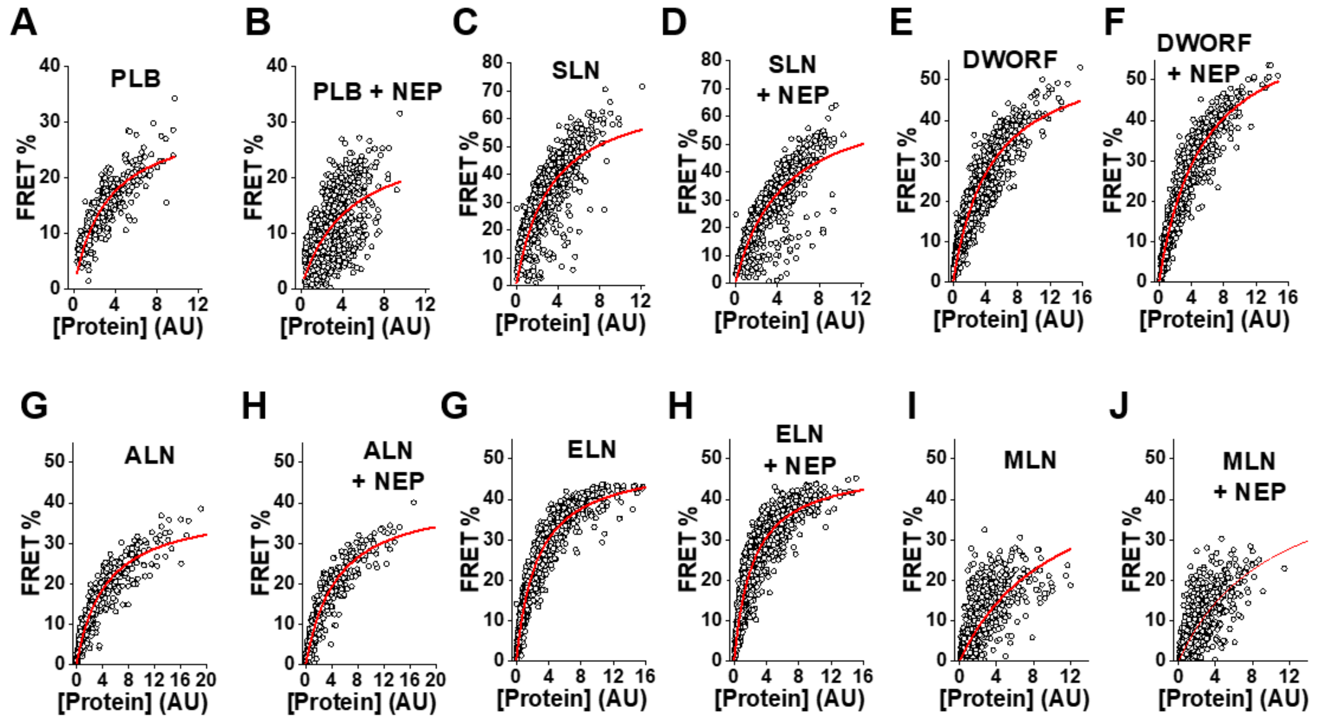


**Supplementary Figure 4.** Representative FRET binding curves between mCer3-SERCA (donor) and YFP-tagged PLB, SLN, ALN, ELN, MLN, or DWORF (acceptors) in HEK293T cells, in the absence or presence of co-expressed NEP. Apparent FRET efficiency is plotted as a function of acceptor fluorescence intensity (a measure of relative expression) and fit to a hyperbolic binding model to obtain the apparent dissociation constant ($K_D$). For ALN, ELN, MLN and DWORF, NEP co-expression produced no significant change in $K_D$.

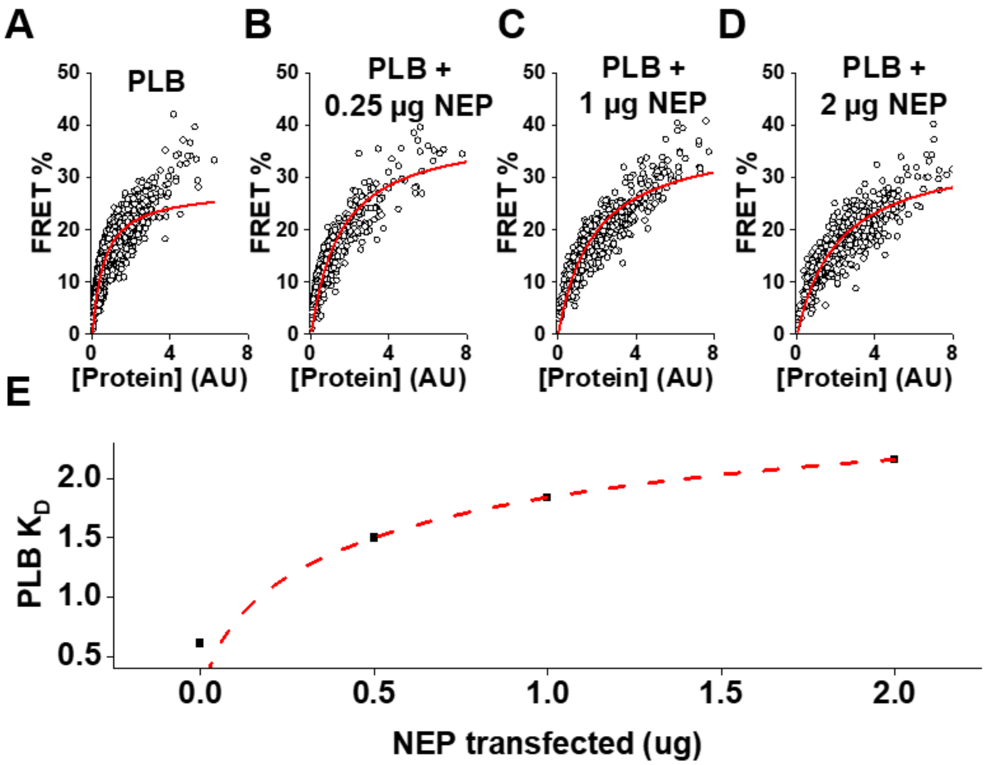


**Supplementary Figure 5. A–E)** FRET binding curves between mCer3-SERCA and YFP-PLB in HEK293T cells co-transfected with increasing amounts of unlabeled NEP plasmid (0, 0.25, 0.5, 1.0, and 2.5 µg per transfection). Apparent K_D values were extracted from hyperbolic fits and the apparent $K_D$'s are plotted in **F)** as a function of NEP plasmid dose. Progressive increases in apparent $K_D$ with increasing NEP expression indicate that NEP-mediated disruption of PLB–SERCA binding is dose-dependent.

| Disease Association | Genes | Count | P-Value |
|---|---|---|---|
| Cardiomyopathy | 1.68% | 23 | 4.74e-3 |
| Fanconi anemia | 0.36% | 5 | 7.36e-2 |

**Supplementary Figure 6.** KEGG pathway enrichment analysis performed on differentially expressed genes from the ET-1 vs control comparison (adjusted $P < 0.05$, $|\log_2 \text{fold change}| \geq 1$) from the bulk RNA-seq dataset in Figure 5. Bars show enriched pathways ranked by adjusted *P*-value, with dot size reflecting gene-set size and color indicating directionality. Pathways related to dilated cardiomyopathy, hypertrophic cardiomyopathy, and adrenergic signaling in cardiomyocytes are significantly enriched, confirming that ET-1 treatment recapitulates transcriptional signatures associated with cardiomyopathy phenotypes.

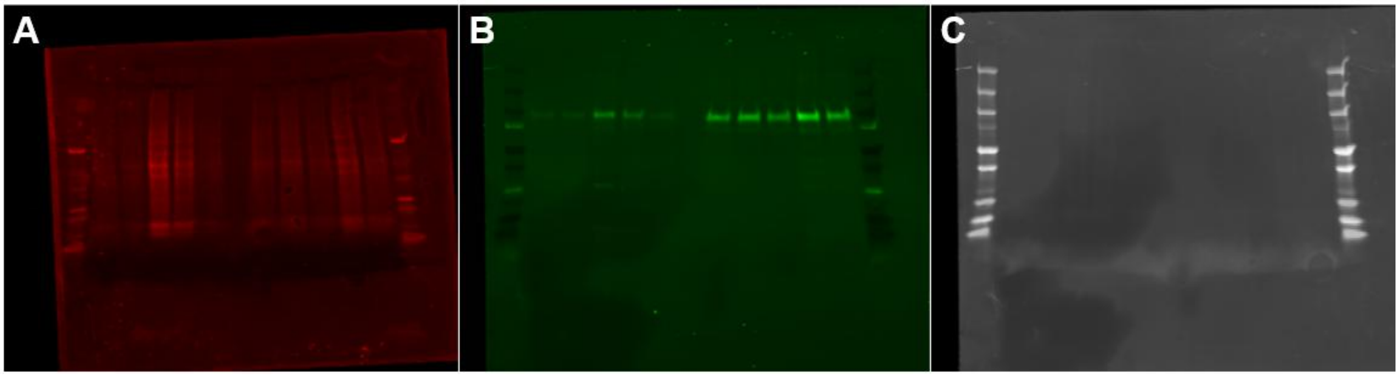


**Uncropped total-protein stain and immunoblot for NEP quantification.** Full, unedited images of the membrane corresponding to the cropped panel in **Figure 1D**. *(A)* Revert total protein stain of the complete membrane, used for lane normalization. *(B)* Anti-CD10 (NEP) immunoblot of the same membrane. Molecular-weight ladder positions (kDa) are indicated at left; the NEP band migrates at ~90 kDa, consistent with the reported mass of mature glycosylated neprilysin [REF/datasheet]. Membrane-enriched left ventricular fractions from non-failing (NF) and dilated cardiomyopathy (DCM) hearts were loaded as in **Figure 1**; lane order matches **Figure 1D–E**.
