## Supplementary material for "Neprilysin mediated cleavage of phospholamban dysregulates SERCA in heart failure": Major Resources Table

In order to allow validation and replication of experiments, all essential research materials listed in the Methods should be included in the Major Resources Table below. Authors are encouraged to use public repositories for protocols, data, code, and other materials and provide persistent identifiers and/or links to repositories when available. Authors may add or delete rows as needed.

### Additional Species Used in This Study (if applicable)

| Strain | Vendor or Source | Background Strain | Sex | Other Information (e.g. breeding scheme for F1, F2…if applicable) |
|---|---|---|---|---|
| Rat (Rattus norvegicus), neonatal, 0–1 day old | Charles River Laboratories | Sprague Dawley (outbred) | Not determined (neonatal pups, pooled) | N/A |

### Antibodies

| Target antigen | Vendor or Source | Catalog # | Working concentration |
|---|---|---|---|
| Neprilysin / CD10 (NEP) | Abcam | ab256494 | IF 1:100; WB 1:1,000 |
| SERCA2 (ATP2A2) | Abcam | ab2817 | IF: 1:500; WB 1:1000 |
| Goat anti-rabbit IgG, AlexaFluor 488 | Abcam | ab150077 | 1:1,000 |
| Goat anti-rabbit/mouse IgG, AlexaFluor 568 | Abcam | ab175471 | 1:1,000 |
| Goat anti-rabbit IgG, IRDye 800CW | LI-COR Biosciences | 926-32211 | 1:10,000 |
| Phalloidin, AlexaFluor 647 (F-actin stain) | Cell Signaling Technology | 8940 | 1:500 |
| DAPI (nuclear stain; in Vectashield) | Vector Laboratories | H-1200 | Stable formula |

### DNA/cDNA Clones

| Clone Name | Sequence | Source / Repository | Persistent ID / URL |
|---|---|---|---|
| Unlabeled human neprilysin (MME) | | Gift from H. Meyer, Universität Osnabrück | Schiemann et al. 2022 (ref 6) |
| mCer3–SERCA2a (human SERCA2a) | | Robia laboratory | GenBank P16615; refs 42, 66, 67 |
| YFP–PLB (human PLN) | | Robia laboratory | UniProt P26678; ref 38 |
| YFP–SLN (sarcolipin) | | Robia laboratory | UniProt O00631; ref 38 |
| YFP–ALN (another-regulin) | | Robia laboratory | UniProt Q8WVX3; ref 38 |
| YFP–ELN (endoregulin) | | Robia laboratory | UniProt P0DI80; ref 38 |
| YFP–MLN (myoregulin) | | Robia laboratory | UniProt P0DMT0; ref 38 |
| YFP–DWORF | | Robia laboratory | UniProt P0DN84; ref 38 |
| YFP–PLB I48X (C-terminal truncation) | | Robia laboratory | Abrol et al. 2014 (ref 16) |

DOI [to be added]

| | | | |
|---|---|---|---|
| YFP–PLB V49A (cleavage-resistant) | | Robia laboratory | ref 16 / this study |
| mCherry–NEP (N-terminal fusion, localization control) | This study | Created using ligation cloning from original construct (Schiemann et al. 2022 (ref 6)) | |
| NEP–mCherry (C-terminal fusion, localization control) | This study | Created using ligation cloning from original construct (Schiemann et al. 2022 (ref 6)) | |

### Cultured Cells

| Name | Vendor or Source | Sex (F, M, or unknown) | Persistent ID / URL |
|---|---|---|---|
| HEK293T | Agilent (Santa Clara, CA) | Female | RRID:CVCL_0063 |
| WTC hiPSC, ACTN2–mEGFP (clone 85) | Allen Cell Collection, via Coriell Institute | Male | AICS-0075-085 |
| Neonatal rat ventricular myocytes (primary) | Isolated in-house (see Animals) | Undetermined | N/A |

### Data & Code Availability

| Description | Source / Repository | Persistent ID / URL |
|---|---|---|
| Fiji / ImageJ (with custom macros) | NIH | RRID:SCR_002285 |
| OriginPro 2025 | OriginLab | RRID:SCR_014212 |
| Proteome Discoverer 2.5 (Sequest HT) | Thermo Fisher Scientific | RRID:SCR_014477 |
| Percolator (PSM validation) | Built into Proteome Discoverer | RRID:SCR_005040 |
| Jalview v2.11.3.2 | Jalview | RRID:SCR_006459 |
| Clustal (alignment, via Jalview web service) | EMBL-EBI | RRID:SCR_001591 |
| GO Functionome | Gene Ontology Consortium | https://functionome.geneontology.org/ |
| MetaMorph | Molecular Devices | RRID:SCR_002368 |
| ZEN Blue (joint deconvolution) | Carl Zeiss | RRID:SCR_013672 |
| NIS-Elements | Nikon | RRID:SCR_014329 |
| IonWizard | IonOptix | Cytosolver3 |
| Xcalibur 4.5 / Tune 3.5 | Thermo Fisher Scientific | RRID:SCR_014593 |

### Other

| Description | Source / Repository | Persistent ID / URL |
|---|---|---|
| Lipofectamine 3000 Transfection Kit | Invitrogen / Thermo Fisher | L3000015 |
| Pierce BCA Protein Assay Kit | Thermo Fisher (Pierce) | 23225 |
| Fura-2 AM | Thermo Fisher | F1221 |
| Revert Total Protein Stain | LI-COR Biosciences | 926-11010 |
| Intercept Blocking Buffer (PBS) | LI-COR Biosciences | 927-70001 |
| Vectashield mounting medium + DAPI | Vector Laboratories | H-1200 |
| Matrigel | Corning | 354277 |
| mTeSR Plus | STEMCELL Technologies | 100-0276 |
| CHIR 99021 | Tocris Bioscience | 4423 |
| IWP 2 | Tocris Bioscience | 3533 |
| B-27 Supplement (± insulin) | Gibco / Thermo Fisher | 17504044 |
| Thiazovivin | STEMCELL Technologies | 72252 |
| Sodium L-lactate | Thermo Scientific Chemicals | L1450014 |
| RPMI 1640, no glucose | Gibco / Thermo Fisher | 11879020 |
| 10X TrypLE | Gibco / Thermo Fisher | A1217701 |

DOI [to be added]

| | | |
|---|---|---|
| KnockOut Serum Replacement | Gibco / Thermo Fisher | 10828028 |
| Saponin | Sigma-Aldrich | 47036 |
| Collagenase + 0.05% trypsin (NRVM digest) | C9891 | C9891 |

| Instrument | Vendor or Source | Model / Part # |
|---|---|---|
| Orbitrap Eclipse Tribrid MS + FAIMS Pro | Thermo Fisher Scientific | — |
| Vanquish Neo UHPLC | Thermo Fisher Scientific | — |
| EASY-Spray analytical column (25 cm) | Thermo Fisher Scientific | ES802A rev2 |
| LSM 880 confocal / Airyscan | Carl Zeiss | — |
| Ti Eclipse 2 / Ti2-E inverted microscope | Nikon | — |
| Prime 95B sCMOS camera | Teledyne Photometrics | — |
| Spectra X light engine | Lumencor | — |
| c600 imaging system | Azure Biosystems | — |
| MyoPacer EP field stimulator + contractility system | IonOptix | — |

### Other — Databases, Deposited Data, and Human Specimens

| Resource / Dataset | Source or Repository | Persistent ID / Accession |
|---|---|---|
| UniProt (Homo sapiens, UP000005640) | UniProt Consortium | RRID:SCR_002380 |
| Bulk RNA-seq dataset (Figure 5) | GEO (NCBI) | GSE335813 |
| Membrane-enrichment proteomics dataset (Figure 1) | ProteomeXchange / PRIDE | *pending* |
| Human LV tissue (NF n=5, DCM n=5; see Suppl. Table 1) | Loyola Cardiovascular Biorepository, Loyola University Chicago (de-identified specimens) | use of de-identified specimens for the present study was determined not to constitute human subjects research |

DOI [to be added]
